# Sphingolipid metabolic flow controls phosphoinositide turnover at the *trans* Golgi network

**DOI:** 10.1101/090142

**Authors:** Serena Capasso, Lucia Sticco, Riccardo Rizzo, Marinella Pirozzi, Domenico Russo, Nina A. Dathan, Felix Campelo, Josse van Galen, Angelika Hausser, Vivek Malhotra, Seetharaman Parashuraman, Alberto Luini, Giovanni D’Angelo

## Abstract

Sphingolipids are membrane lipids, which are globally required for eukaryotic life. Sphingolipid composition varies among endomembranes with pre- and post-Golgi compartments being poor and rich in sphingolipids, respectively. Thanks to this different sphingolipid content, pre- and post-Golgi membranes serve different cellular functions. Nevertheless, how subcellular sphingolipid levels are maintained in spite of trafficking and metabolic fluxes is only partially understood. Here we describe a homeostatic control circuit that controls sphingolipid levels at the trans Golgi network. Specifically, we show that sphingomyelin production at the trans Golgi network triggers a signalling reaction leading to PtdIns(4)P dephosphorylation. Since PtdIns(4)P is required for cholesterol, and sphingolipid transport to the trans Golgi network, PtdIns(4)P consumption leads to the interruption of this transport in response to excessive sphingomyelin production. Based on this evidence we envisage a model where this homeostatic circuit maintains the sphingolipid composition of trans Golgi network and thus of post-Golgi compartments constant, against instant fluctuations in the sphingolipid biosynthetic flow.

## Introduction

Sphingolipids (SLs) are bioactive components of eukaryotic membranes (Hannun & Obeid, 2008). Several stimuli activate SL metabolic enzymes to produce SL species that act as first or second messengers in signalling programmes involved in irreversible cell fate decisions (Hannun & Obeid, 2008) and where their derangements often result in the development of cancer (Morad & Cabot, 2013). Not surprisingly then, SL metabolism is subjected to control modules that maintain its balance and prevent unintended drifts away from the steady state (Breslow & Weissman, 2010, Cantalupo, Zhang et al., 2015, Siow & Wattenberg, 2012, Vacaru, Tafesse et al., 2009). While these controls account for global SL homeostasis, SL biosynthesis involves the regional production and intracellular transport of different metabolites. As a consequence SLs are unevenly distributed among cell membranes with *post*-Golgi membranes being rich and *pre*-Golgi membranes being poor in SLs (Holthuis & Menon, 2014, van Meer, Voelker et al., 2008). Importantly, high concentrations of SLs and cholesterol at *post*-Golgi membranes increase membrane thickness and promote the formation of membrane domains, both of which influence protein trafficking, and signalling and thus affect cellular functions (Hannun & Obeid, 2008, Holthuis & Menon, 2014, Lingwood & Simons, 2010, Patterson, Hirschberg et al., 2008, Sharpe, Stevens et al., 2010, van Galen, Campelo et al., 2014).

A hub in establishing the intracellular SLs distribution is the Golgi complex (De Matteis & Luini, 2008, Hannun & Obeid, 2008, Holthuis & Menon, 2014). In fact, newly synthesized ceramide (Cer) in the endoplasmic reticulum (ER) is either transported by vesicular intermediates to the *cis*-Golgi pole, where it is converted to glucosylceramide (GlcCer), or delivered to the *trans* Golgi network (TGN) by the Cer transfer protein (CERT), for the production of sphingomyelin (SM) (Hanada, Kumagai et al., 2003). Similarly, *cis*-Golgi produced GlcCer can progress through Golgi *cisternae* being converted to ganglio series glyco-SLs (GSLs) or shunted to the TGN by the GlcCer transfer protein FAPP2 (D‘Angelo, Polishchuk et al., 2007, D‘Angelo, Uemura et al., 2013) where it is used to produce globoseries GSLs (D‘Angelo et al., 2013) (**Fig EV1**).

CERT and FAPP2 are recruited to the TGN by the phosphoinositide phosphatidylinositol-4-phosphate [PtdIns(4)*P*] (D‘Angelo, Vicinanza et al., 2008, Yamaji & Hanada, 2015). As a consequence, PtdIns(4)*P* is required for both SM and GSLs syntheses and enrichment at *post-*Golgi membranes (D‘Angelo et al., 2013, Toth, Balla et al., 2006). PtdIns(4)*P* levels at the TGN depend on its production (by TGN-localized PtdIns4-kinases) and consumption (by the ER-localized PtdIns-4-phosphatase Sac1) (De Matteis, Wilson et al., 2013). The lipid transfer protein OSBP1 (Mesmin, Bigay et al., 2013) operates the transport of PtdIns(4)*P* from the TGN to the ER for its dephosphorylation by Sac1. This trafficking/metabolic step is accomplished at specific sites of close apposition between ER and TGN defined as ER-TGN membrane contact sites (MCSs) where it is coupled to the transport of cholesterol from the ER to the TGN (Mesmin et al., 2013).

Once at the TGN, SLs, GSLs, and cholesterol are incorporated in membrane carriers and transported to their *post*-Golgi destinations thus contributing to the establishment of the *post-*Golgi membrane territory (De Matteis & Luini, 2008, Deng, Rivera-Molina et al., 2016, Klemm, Ejsing et al., 2009). Nevertheless, carriers formation at the TGN occurs continuously so that TGN membranes are subjected to constant dissipation and rebuilding according to secretory flux volume to the TGN (De Matteis & Luini, 2008). Moreover, fluctuations in SL *de novo* synthesis are reported under a variety of signalling and stress conditions (Hannun & Obeid, 2008). Thus it is not fully understood how cells keep the local TGN lipid composition (and as a consequence that of *post*-Golgi membranes) controlled in spite of uncoordinated changes in membrane trafficking and SL precursors supply.

Here we have addressed this issue by acutely modulating the SL flow to the Golgi complex to then measure the effect on TGN composition and metabolic capacity. Our results indicate that the SL flow controls the PtdIns(4)*P* levels at the TGN. Specifically we describe a SL dependent signalling leading to PtdIns(4)*P* consumption and consequent release of PtdIns(4)*P* binding proteins from TGN membranes. Provided that PtdIns(4)*P* is required for SL and cholesterol transport to the TGN (D‘Angelo et al., 2013, Toth et al., 2006), our findings reveal a negative feedback circuit that ensures the homeostatic control of SL levels at the TGN and at *post-*Golgi membranes.

## Results

### Adaptive metabolic response to SL flow

To investigate how cells respond to acute changes in SL flow, HeLa cells were subjected to controlled perturbations in the SL metabolic input. Cells were first treated with Fumonisin B1, an inhibitor of Cer synthesis and SL recycling (Ogretmen, Pettus et al., 2002, Wang, Norred et al., 1991) (FB1; 50μM for 24hours) or myriocin, a serine-palmitoyl transferase inhibitor (Horvath, Sutterlin et al., 1994) (2.5μM for 24 hours) to obliterate SL production (**Fig EV1**). Subsequently, cells were fed for 2 hours with increasing concentrations of cell permeable SL precursors N-hexanoyl-D-*erythro*-sphingosine (C6-DCer; for FB1 treated cells) or D-*erythro*-sphingosine (D-Sph; for myriocin treated cells) and the amount of SL precursor converted either to SM or GlcCer was quantified. As shown in **Fig 1A** the fractions of SL precursors converted to SM or GlcCer were constant over a wide range (from 5nM to 1μM) of C6-D-Cer and D-Sph concentrations. In contrast addition of higher amounts of SL precursors resulted in a sudden decrease of SL flux directed to SM synthesis, which was counterbalanced by an increase in GlcCer synthesis and Cer accumulation (**Fig 1A**).

**Figure 1:**
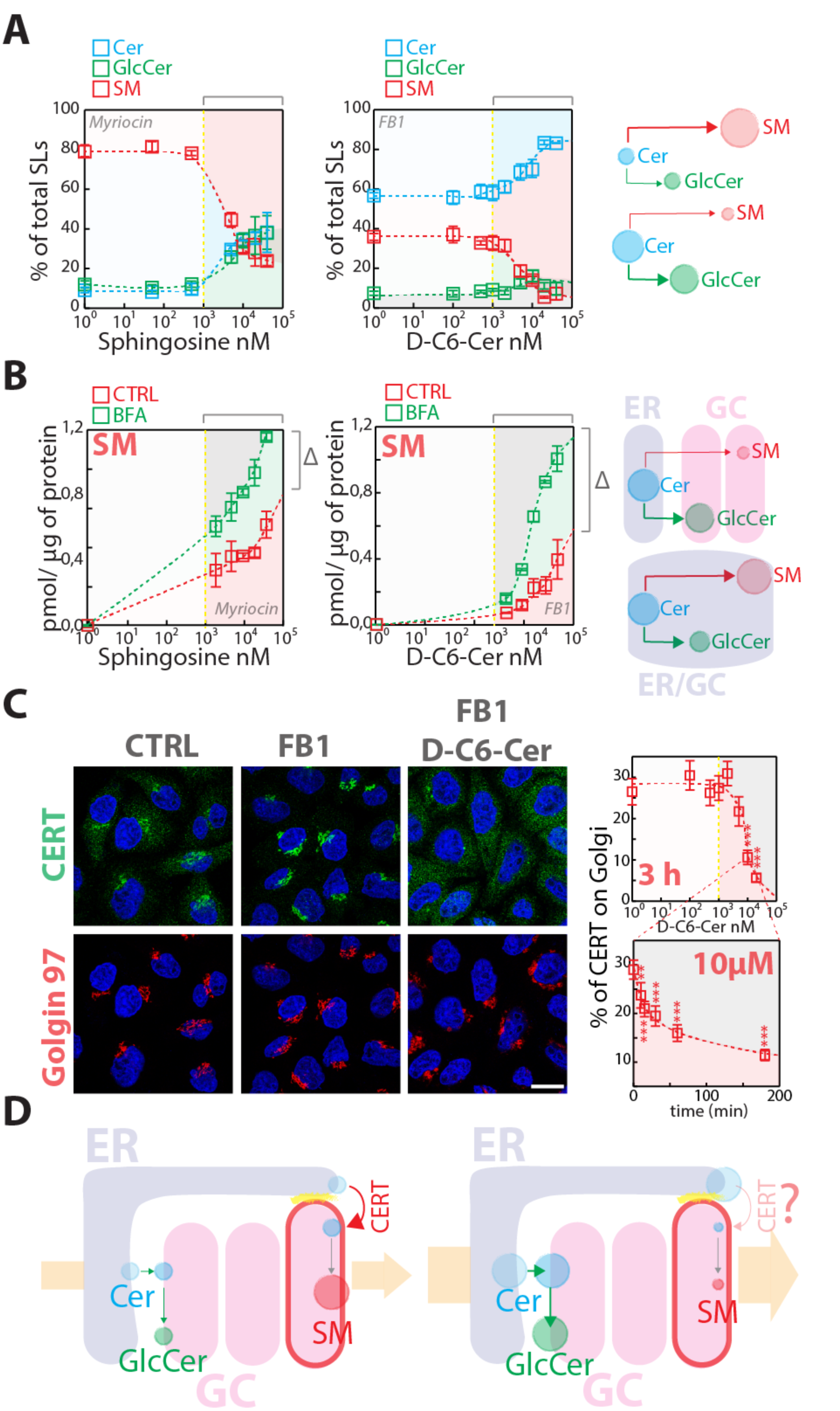
Metabolic cell response to SL flow. **A)** HeLa cells treated with myriocin (2.5 μM) or FB1 (50μM) for 24 hours (left and middle panel respectively) were fed with increasing concentrations of D-Sph (left panel) or C6-DCer (middle panel) for 2 hours. ^3^H-D-Sph or ^3^H-C6-D-Cer (≈5nM) were mixed as tracers with their non-radioactive counterparts. The percentage of total radioactivity associated with Cer (cyan), GlcCer (green) and SM (red) in the different conditions was quantitated after lipid extraction and HPTLC separation. The right panel schematizes the metabolic effect of sustained SL flow. **B)** (Left and middle panels). Cells treated as in **(A)** were subjected to Brefeldin A (BFA) administration (5μg/mL) before (30 min pre-treatment) and during SL administration for 2 hours. The amount of SM (pmol) produced under control (red squares) or BFA (green squares) per μg of protein lysate was calculated as detailed in **Methods** and indicated at increasing SL precursor concentrations. Δ indicates the difference in SM production at saturating SL precursors amounts. The right panel schematizes the effect of BFA on SM production. **C)** CERT localisation in HeLa cells control (CTRL), treated with FB1 (50μM, 24 hours), or with 10μM C6-D-Cer for 2 hours after 24 hours treatment with FB1. Bar, 10 μm. Left panels indicate the percentage of CERT associated with the Golgi complex at increasing C6-D-Cer concentrations (upper graph) and at the indicated times (lower graph). Data are means ± S.E.M. of at least 3 independent experiments. **D)** Schematic representation of SL flow induced redistribution of CERT to the cytosol and consequent inhibition of SM synthesis.

The fact that SM synthesis is impaired at high SL precursor concentrations can be due either to saturation of enzymes synthesizing SM (i.e., mostly SM synthase 1 [SMS1]) (Huitema, van den Dikkenberg et al., 2004) or of the system that supplies Cer to SMS1 (i.e., CERT (Hanada et al., 2003)). To discriminate between these possibilities we repeated the experiment described in **Fig 1A** in cells pre-treated with Brefeldin A (BFA; 5μg/mL). BFA redistributes Golgi membranes to the ER (Lippincott-Schwartz, Yuan et al., 1989) and relocates SMS1 and GlcCer synthase (GCS) to this mixed ER-Golgi compartment (**Fig EV2**). Under these conditions Cer is readily available to both GCS and SMS1 with no need for either vesicular or non-vesicular Cer transport (D‘Angelo et al., 2013). As shown in **Fig 1B** BFA treatment resulted in increased SM synthesis at high precursor concentrations (> 1μM), suggesting that the observed drop in SM synthesis (**Fig 1A**) is not due to saturation of SMS1.

We then analysed CERT localization in response to SL perturbations. As already reported by others (Kumagai, Kawano et al., 2007) we found that SL depletion by either FB1 or myriocin induced CERT recruitment to the TGN (**Fig 1C**, **Fig EV3**, and **4**). Interestingly, when cells were treated with C6-D-Cer or D-Sph, CERT shifted to a cytosolic distribution at concentrations similar to those inducing a drop in SM production (**Fig 1C,** and **Fig EV3**).

Moreover, CERT delocalization was complete within 60 min of treatment (**Fig 1C**). Based on this evidence we concluded that SL flow acutely counteracts CERT recruitment to the TGN (and as a consequence SM synthesis) (**Fig 1D**). We thus wondered what mechanism could be responsible for this phenomenon.

### Sustained SL flow induces the consumption of the TGN pool of PtdIns(4)P

First we looked for peripheral Golgi proteins, which behaved similarly to CERT in terms of association with membranes in response to sustained SL flow. While most of the tested proteins remained associated with the Golgi membranes even after 2 hours of C6-D-Cer treatment, a specific subset of TGN proteins was displaced from the Golgi membranes. This subset contained the proteins FAPP2, OSBP1, GOLPH3, and AP-1 (γ-adaptin) (**Fig 2A** and **Fig EV5**).

**Figure 2:**
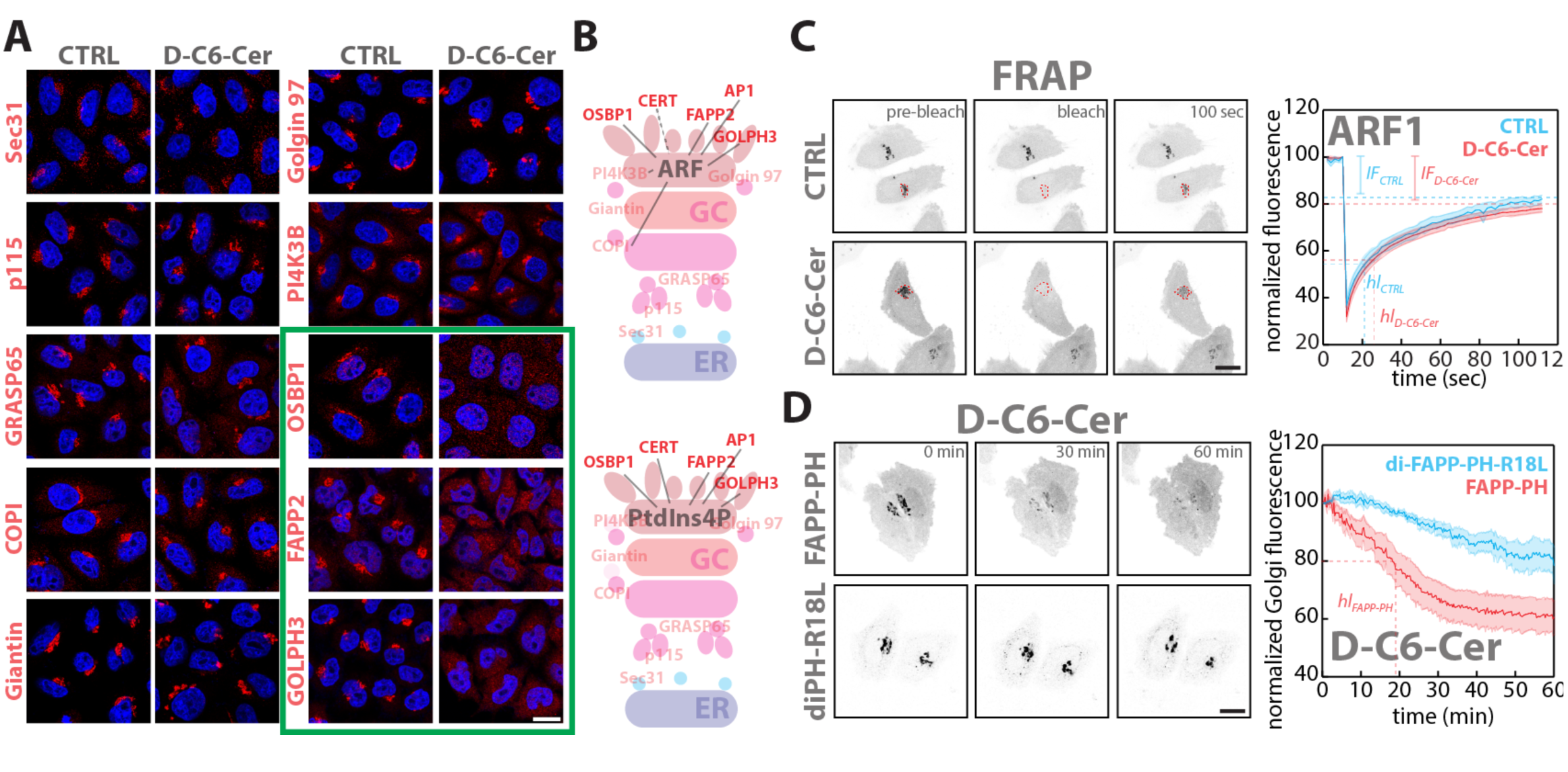
The localization of TGN PtdIns(4)P effectors is sensitive to SL flow. **A)** Cells were treated with either vehicle (EtOH) or D-C6-Cer (10μM) for 2 hours, fixed and stained for nuclei (DAPI; blue) and with antibodies to different Golgi associated proteins (red). **B)** Schematic representation of Golgi-proteins localization. Upper panel shows proteins that require ARF and lower panel proteins that require PtdIns(4)*P* for their Golgi localization. In solid red are proteins sensitive to sustained SL flow. **C)** FRAP based assessment of ARF1-GFP dynamics of association/ dissociation from Golgi membranes in EtOH or D-C6-Cer (10μM) treated cells (see **Methods** for details; left panels). Mean normalized fluorescence intensity ± SEM over time under CTRL (cyan) and D-C6-Cer (red) treatment is reported (*hl*: half life; *IF*: immobile fraction). **D)** Live imaging analysis (1 hour) of FAPP-PH-GFP or diFAPP-PH-R18L-GFP during D-C6-Cer treatment (10 μM). Mean normalized fluorescence intensity ± SEM over time is reported for FAPP-PH-GFP (red) and diFAPP-PH-R18L-GFP (cyan) (*hl*: half life); Bars, 10μm.

Given that these proteins require ARF and PtdIns(4)*P* for their recruitment to the TGN (**Fig 2B**) (De Matteis et al., 2013) the effect of SL flow on ARF1 and PtdIns(4)*P* was investigated. As shown in **Fig EV6,** C6-D-Cer treatment did not perturb ARF1 localization to the Golgi membranes. Moreover, when the dynamics of ARF1-GFP association to the Golgi complex were examined by Fluorescence Recovery After Photobleaching (FRAP) experiments in cells treated with C6-D-Cer (10μM for 2 hours) (**Fig 2C**; **Movie 1** and **2**) no differences were observed.

Next, the effect of C6-D-Cer on PtdIns(4)*P* was determined. For this purpose we initially used the GFP-tagged Pleckstrin Homology domain of FAPP2 (FAPP-PH-GFP) as a PtdIns(4)*P* probe (D‘Angelo et al., 2013, Dowler, Currie et al., 2000, Godi, Di Campli et al., 2004). As shown in **Fig 2C** and in **Movie 3**, C6-D-Cer administration (10μM) induces the redistribution of FAPP-PH from the Golgi complex to the cytosol within 60 min. On the other hand, a tandem FAPP-PH mutant (diFAPP-PH-R18L-GFP), which localizes to the Golgi complex in spite of being unable to bind PtdIns(4)*P* (Godi et al., 2004) was less sensitive to this treatment (**Fig 2C** and in **Movie 4**), suggesting that SL flow to the Golgi complex controls the TGN phosphoinositide composition and not ARF1 localization.

Indeed both C6-D-Cer and D-Sph treatments induced the disappearance of the PtdIns(4)*P* signal from the Golgi region (**Fig 3A** and **Fig EV7**) as assessed by the use of anti-PtdIns(4)*P* antibody (Hammond, Schiavo et al., 2009). Intriguingly, the C6-D-Cer concentration and times of treatment causing PtdIns(4)*P* loss were superimposable to those that induce CERT relocation to the cytosol and thus its inactivation (compare **Fig 1C** and **3B)**.

**Figure 3:**
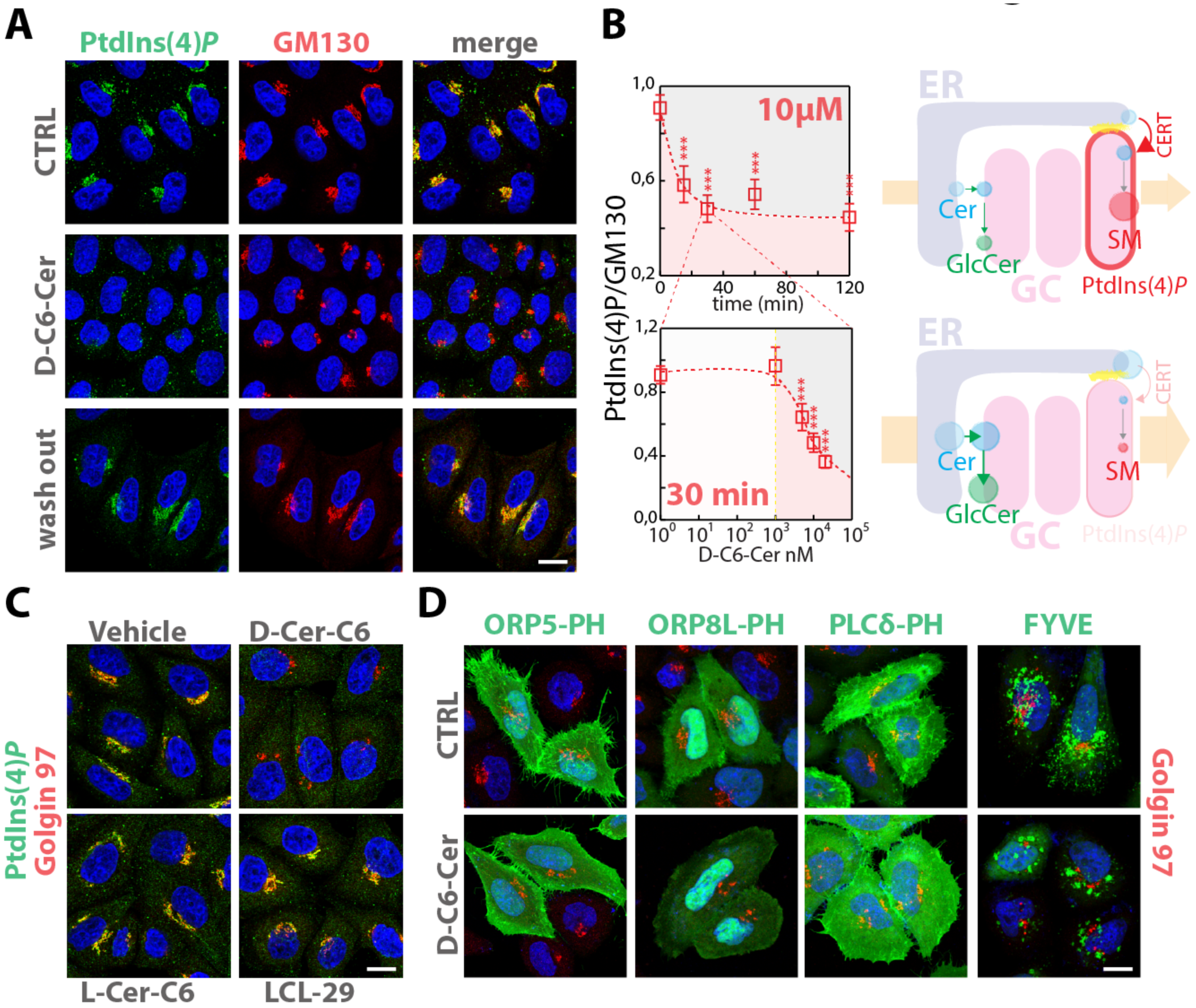
SL flow controls PtdIns(4)P levels at the Golgi. **A)** Cells treated either with vehicle (EtOH), D-C6-Cer (10μM) for 30 min, or treated with DC6-Cer (10μM) for 30 min and washed out for 4 hours were stained with a specific anti PtdIns(4)*P*-antibody as detailed in **Methods**. **B)** HeLa cells treated with D-C6-Cer (10μM) for the indicated times (upper left panel) or with increasing D-C6-Cer concentrations for 30 min (lower left panel) were processed and stained as in **(A)**. Confocal images were acquired, segmented, and analysed by CellProfiler software, as detailed in **Methods**. Average of normalized PtdIns(4)*P* / GM130 integrated intensities ratio is reported for each time or concentration point. Data are means ± 5 x S.E.M of values obtained from at least 500 cells per experimental point. Right panels schematize the effect of SL flow on PtdIns(4)*P* levels, CERT recruitment to the Golgi, and SM synthesis. **C)** HeLa cells treated either with vehicle (EtOH), D-C6-Cer (10μM), L-C6-Cer (10μM), or LCL-29 (10 μM) for 30 min were fixed and permeabilized and stained as in **(A)**. **D)** Cells transfected with GFP-tagged ORP5-PH, ORP8L-PH, PLCδ-PH, or FYVE were treated either with vehicle (EtOH), D-C6-Cer (10μM) for 2 hours, fixed and stained with DAPI (blue), and an anti-Golgin97 antibody (red). Bars, 10 μm.

While PtdIns(4)*P* loss (**Fig 3B**) is in good agreement with the effects measured on SM synthesis inhibition by C6-D-Cer (**Fig 1A**), it also predicts an inhibitory effect of C6-D-Cer treatment on globo-series GSL production as this depends on FAPP2 and its ability to bind PtdIns(4)*P* at the TGN (D‘Angelo et al, 2013b). Accordingly a strong inhibitory effect on globo-series GSLs (Gb3) synthesis (≈83% reduction) was measured in cells treated with D-C6-Cer (10μM, 2h), which was counterbalanced by a significant accumulation of GlcCer (≈56% increase) (**Fig EV8**).

To exclude that the effects observed under SL challenges were caused by non-specific cell poisoning control experiments were performed. According to these experiments: **(i)** the effect of C6-D-Cer and D-Sph is reversible as cells reacquire normal PtdIns(4)*P* staining after wash-outs (**Fig 3A**, and **Fig EV7**). **(ii)** When the non-metabolizable C6-L-Cer enantiomer (Duran, Campelo et al., 2012) was added to the cells no effect was observed on PtdIns(4)*P* staining (**Fig 3C**). **(iii)** Since Cer exerts cytostatic effects (Hannun & Obeid, 2008) and growth conditions influence PtdIns(4)*P* levels at the TGN (Bajaj Pahuja, Wang et al., 2015, Blagoveshchenskaya, Cheong et al., 2008), the possibility that the observed effect results from an indirect effect of Cer on cell proliferative signalling was considered. Thus cells were treated with a Cer analogue D-erythro-2-N-[6‘-(1‘‘pyridinium)hexanoyl]-sphingosine bromide (LCL29) that inhibits cell growth without being concentrated and metabolized at the Golgi complex (Hou, Jin et al., 2011). Under these conditions we observed no changes in PtdIns(4)*P* staining at the Golgi (**Fig 3C**), suggesting that Cer needs to be transported and metabolized at the Golgi complex to influence PtdIns(4)*P* levels.

In addition to the TGN, PtdIns(4)*P* pool is also enriched at the plasma membrane, where it is recognized by the PH domains of OSBP family proteins ORP5 and ORP8L (Chung, Torta et al., 2015). As shown in **Fig 3D** C6-D-Cer did not influence the plasma membrane association of GFP tagged versions of ORP5 and ORP8L PH domains. Similarly, C6-DCer treatment did not induce appreciable changes in the membrane association of probes for other phosphoinositides (i.e., GFP-PH-PLCδ and GFP-FYVE, recognizing PtdIns(4,5)*P*_2_, and PtdIns(3)*P* respectively (Burd & Emr, 1998, Lemmon, Ferguson et al., 1995, Stauffer, Ahn et al., 1998, Varnai & Balla, 1998); **Fig 3D**). This indicates that the TGN pool of PtdIns(4)*P* is specifically sensitive to sustained SL flow with consequences on local SM and GSLs production.

### SM synthesis induces the consumption of PtdIns(4)P at the TGN

According to the data so far reported Cer requires to be transported to the Golgi complex to induce its effect on TGN PtdIns(4)*P*. At the Golgi, Cer is metabolized to GlcCer by GCS (Ichikawa, Ozawa et al., 1998) or to SM by SMS1 (Huitema et al., 2004). To investigate the specific role of these reactions in PtdIns(4)*P* consumption, GCS1 and SMS1 were silenced and cells were treated with D-C6-cer for 2 hours. As shown in **Fig 4A** SMS1-KD specifically protected TGN PtdIns(4)*P* from C6-D-Cer treatment. As a consequence of this effect, SMS1-KD prevents PtdIns(4)*P* binding proteins (here we show GOLPH3) from being redistributed to the cytosol after C6-D-Cer treatment (**Fig EV9A**). Interestingly, untreated SMS1-KD cells showed increased association of CERT, OSBP1, and GOLPH3-GFP to the TGN (**Fig EV9B, C**), possibly due to increased local PtdIns(4)*P* levels upon reduced basal SM synthesis. Along similar lines, silencing of CERT, which mediates Cer transport to the TGN for SM synthesis (Hanada et al., 2003), but not of FAPP2, which mediates GlcCer transport to the TGN for globo-series GSL synthesis (D‘Angelo et al, 2013b), also protected Golgi PtdIns(4)*P* from C6-D-Cer treatment (**Fig EV10**).

**Figure 4:**
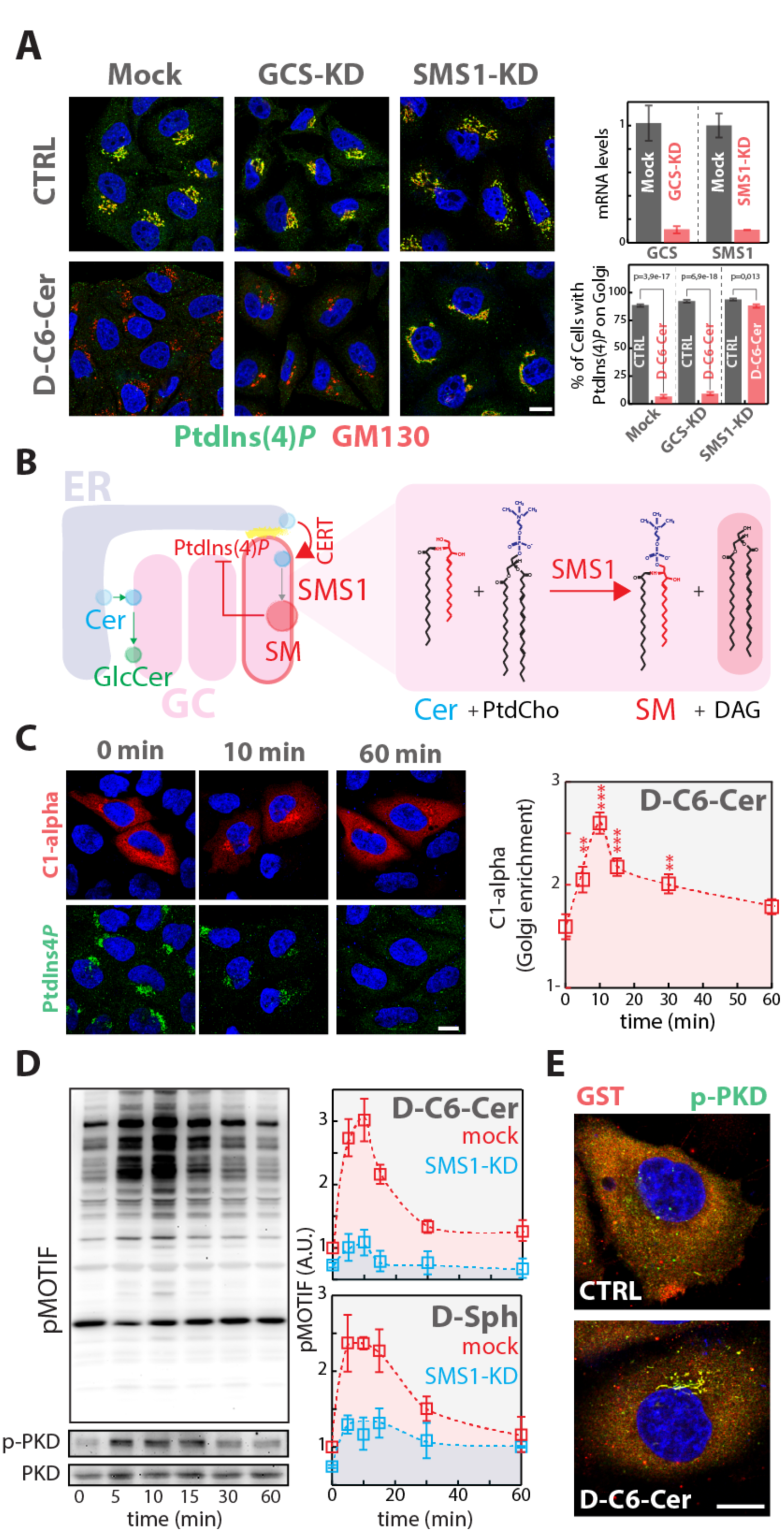
Cer to SM conversion controls PtdIns(4)P levels at the TGN. **A)** Cells KD for GCS or SMS1 were treated with D-C6-Cer (10μM) for 2 hours, fixed, permeabilized, and stained as in **Figure 3A**. KD efficiencies are indicated as reduction in mRNA levels as assessed by RT qPCR (upper right panel). The percentage of cells showing PtdIns(4)*P* staining at the Golgi in CTRL and D-C6-Cer treated cells is reported (lower right panel). **B)** Schematic representation of the feedback loop regulating SM synthesis through PtdIns(4)*P* (left panel). Biochemical depiction of Cer to SM conversion operated by SMS1. **C)** Cells transfected expressing the GST-tagged DAG sensor PKDC1α domain (C1-alpha) were treated for the indicated times with D-C6-Cer (10 μM), fixed, permeabilized and stained with DAPI (blue), anti-GST (red), anti- PtdIns4*P* (green) antibodies (left panel). The Golgi associated C1-alpha fraction at the different time-points is reported (right panel). **D)** Cells treated with D-C6-Cer (10μM) for the indicated times were lysed and lysates were processed for SDS-PAGE and Western Blotting (as detailed in methods). Antibodies recognizing total PKD (PKD), phospho-Ser916-PKD (p-PKD), and phosphorylated PKD substrates (pMOTIF) were used to monitor PKD activation (left panels). PKD activation in both mock (red) and SMS1-KD (cyan) cells was evaluated by quantitating phosphorylation of PKD substrates during D-C6-Cer (10μM) or D-Sph (30μM) administration. **E)** Cells transfected with a plasmid encoding GST-PKD2 were treated with D-C6-Cer (10μM) for 15 minutes, fixed, permeabilized and stained with DAPI (blue), anti-GST (red), and anti-p-PKD (green). Bars, 10μm. Data are means ± S.D of at least 3 independent experiments

The SMS1-mediated conversion of Cer to SM is coupled to the production of diacylglycerol (DAG) (**Fig 4B**) (Hannun & Obeid, 2008). Thus DAG production at the TGN was investigated in relation to PtdIns(4)*P* loss in response to SL flow. HeLa cells were, thus, transfected with a vector encoding the C1α domain of PKD (PKD-C1α), which is recruited to membranes by DAG (Baron & Malhotra, 2002). PKD-C1α localization was then followed during C6-D-Cer administration along with that of PtdIns(4)*P*. As shown in **Fig 4C**, C6-DCer treatment induced a fast recruitment of PKD-C1α to the Golgi region that peaked at 10-15 min and that preceded the disappearance of PtdIns(4)*P*. Interestingly, PKD-C1α recruitment dropped sharply at 30-60 min of treatment coinciding with the loss of a detectable PtdIns(4)*P* signal, which suggests that C6-D-Cer administration induces a transient burst in DAG production at the TGN before PtdIns(4)*P* is consumed.

Since DAG at the TGN induces PKD recruitment and activation (Baron & Malhotra, 2002), we treated cells with C6-D-Cer for different times and monitored PKD autophosphorylation (indicative of activation) by Western Blotting and immunofluorescence. As shown in **Fig 4D** C6-D-Cer treatment induced PKD activation that peaked at 10-15 min treatment, followed by a decrease of PKD phosphorylation at 30 and 60 min. As a consequence of this dynamics also PKD substrates (as recognized by a specific anti-PKD phospho-motif; [pMOTIF] (Doppler, Storz et al., 2005) were phosphorylated at early time points of C6-D-Cer treatment, to then be dephosphorylated after 30 and 60 min of treatment (**Fig 4D**). The phosphorylation of PKD substrates induced by C6-D-Cer was largely dependent on Cer to SM conversion as SMS1-KD inhibited their response to C6-DCer treatment. Similar results were obtained by D-Sph, which also induced a transient and SMS1-dependent PKD substrate phosphorylation (**Fig 4D**). When the subcellular location of PKD activation upon C6-D-Cer treatment was visualised by immunofluorescence, a strong activation signal in the perinuclear region was noticed (**Fig 4E**).

Given the sharp decrease in PKD activity observed after 30 min of C6-D-Cer and D-Sph treatments, we wondered whether this was a consequence of reduced DAG at the TGN or if these treatments would render PKD irresponsive to stimulatory inputs. To test this, cells were treated for 15 minutes with the DAG analogue Phorbol 12,13-diButyrate which activates PKD (Chen, Deng et al., 2008) before or after 30 min of C6-D-Cer treatment. As shown in **Fig EV11A**, PdBu induced PKD activation irrespective of C6-D-Cer treatment, suggesting that PKD is still responsive after C6-D-Cer treatment. As PtdIns(4)*P* (PdBu; 1μM) is required for Cer transport to the TGN and for SM (and thus DAG) production (Hanada et al., 2003, Toth et al., 2006), we reasoned that the PtdIns(4)*P* drop and the consequent inhibition of SM production induced by C6-D-Cer are the cause of the reduced PKD activity.

To test this hypothesis cells were treated with the PI4K inhibitor PIK93 (Toth et al., 2006) before C6-D-Cer administration and PKD activation was evaluated by measuring the phosphorylation of its substrates. As shown in **Fig EV11B,** PIK93 treatment rendered cells unresponsive to C6-D-Cer in terms of PKD activation, supporting the hypothesis that PtdIns(4)*P* is required for SL induced PKD activation and that its consumption terminates PKD signalling.

Given the association between PKD signalling and PtdIns(4)*P* consumption induced by SM production we wondered whether PKD itself could be involved in the control of PtdIns(4)*P* levels at the TGN. We thus overexpressed a wt or dominant negative form of PKD (PKD-KinDead, PKD-K618N) (Liljedahl, Maeda et al., 2001) in HeLa cells before C6-D-Cer treatment. As shown in **Fig 5A** PKD-KinDead expression protected PtdIns(4)*P* from C6-D-Cer induced consumption while PKD-wt did not. Along similar lines the knockdown of all three PKD isoforms (Rykx, De Kimpe et al., 2003) induced a substantial protection towards C6-D-Cer treatment (**Fig 5B**), indicating that transient PKD activation is required for the consumption of the TGN PtdIns(4)*P* pool under C6-D-Cer treatment. When the individual contribution of each isoform to this phenotype was considered, PKD2 was found to be responsible for C6-D-Cer induced PtdIns(4)*P* consumption (**Fig 5C**). Altogether these data indicate that SM synthesis, through PKD activation, controls PtdIns(4)*P* levels at the TGN.

**Figure 5:**
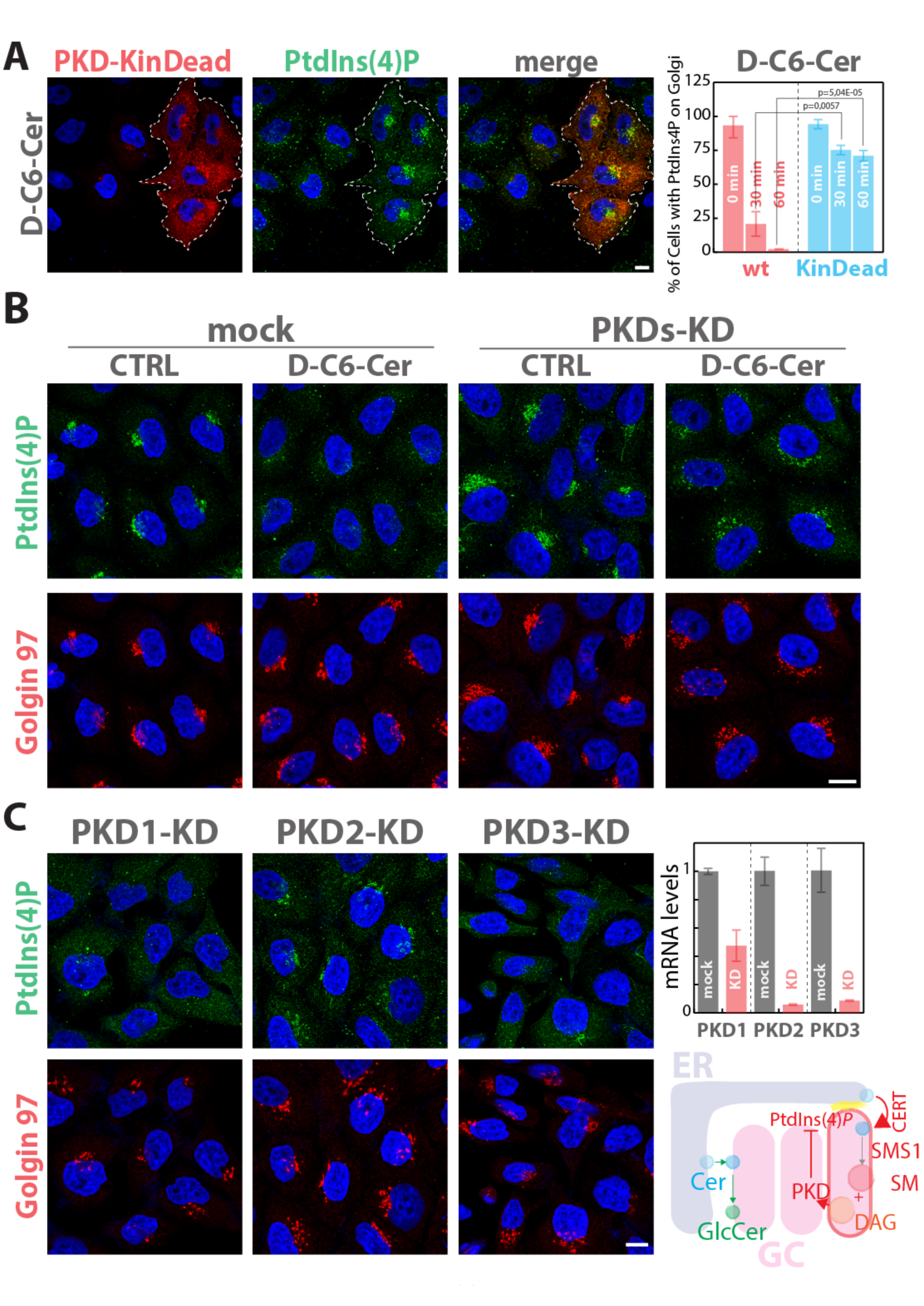
PKD activation is required for SL flow induced PtdIns(4)P consumption. **A)** Cells expressing GST-tagged versions of either PKD wild type (PKD-wt) or of a kinase dead mutant of PKD (PKD-KinDead; PKD-K618N) (Liljedahl et al., 2001) were treated for 30 and 60 minutes with D-C6-Cer (10μM). After fixation and permeabilization cells were stained with DAPI (blue), anti GST (red), and anti-PtdIns(4)*P* (green). The left panel shows a representative image of PKD-KinDead transfected cells after 60 min of D-C6-Cer (10μM) treatment. The percentage of PKD transfected (wt, red; KinDead, cyan) cells showing PtdIns(4)*P* staining at the Golgi at the different times of D-C6-Cer treatment is reported (right panel). **B)** Cells silenced for the three PKD isoforms were treated with D-C6-Cer (10μM) for 60 min, fixed, permeabilized, and stained as in **Figure 3A**. **C)** Cells silenced for each of the three PKD isoforms were treated with D-C6-Cer (10μM) for 60 min, fixed, permeabilized, and stained as in **Figure 3A**. KD efficiency for each PKD isoform is reported (mRNA levels as measured by RT qPCR; upper right panel). Schematic representation of SL mediated PKD activation and PKD dependent PtdIns(4)*P* consumption at the TGN (lower right panel).

### SL flow regulates phosphoinositide turnover through PI4KIIIβ and Sac1

PtdIns(4)*P* is synthesized through the phosphorylation of PtdIns by PtdIns-4-kinases (PI4KIIα, PI4KIIIα, PI4KIIβ, and PI4KIIIβ), while the opposite reaction is catalysed by the PtdIns(4)*P*-4-phosphatase Sac1 (De Matteis et al., 2013) (**Fig 6A**). We, thus, wondered which of these enzymes is involved in the SL mediated modulation of PtdIns(4)*P* pool at the TGN. To this aim we individually silenced each PI4K or Sac1 in HeLa cells and treated them with myriocin or C6-D-Cer to inhibit or stimulate the SL metabolism. We then evaluated both myriocin-induced CERT recruitment to the Golgi complex and C6-D-Cer triggered PtdIns(4)*P* consumption. As shown in **Fig 6B** PI4KIIIβ-KD specifically abolished CERT recruitment upon myriocin treatment and PtdIns(4)*P* staining at the TGN in both vehicle and C6-D-Cer treated cells, suggesting that PI4KIIIβ is the main PI4K involved in the production of the PtdIns(4)*P* pool regulated by SLs.

**Figure 6:**
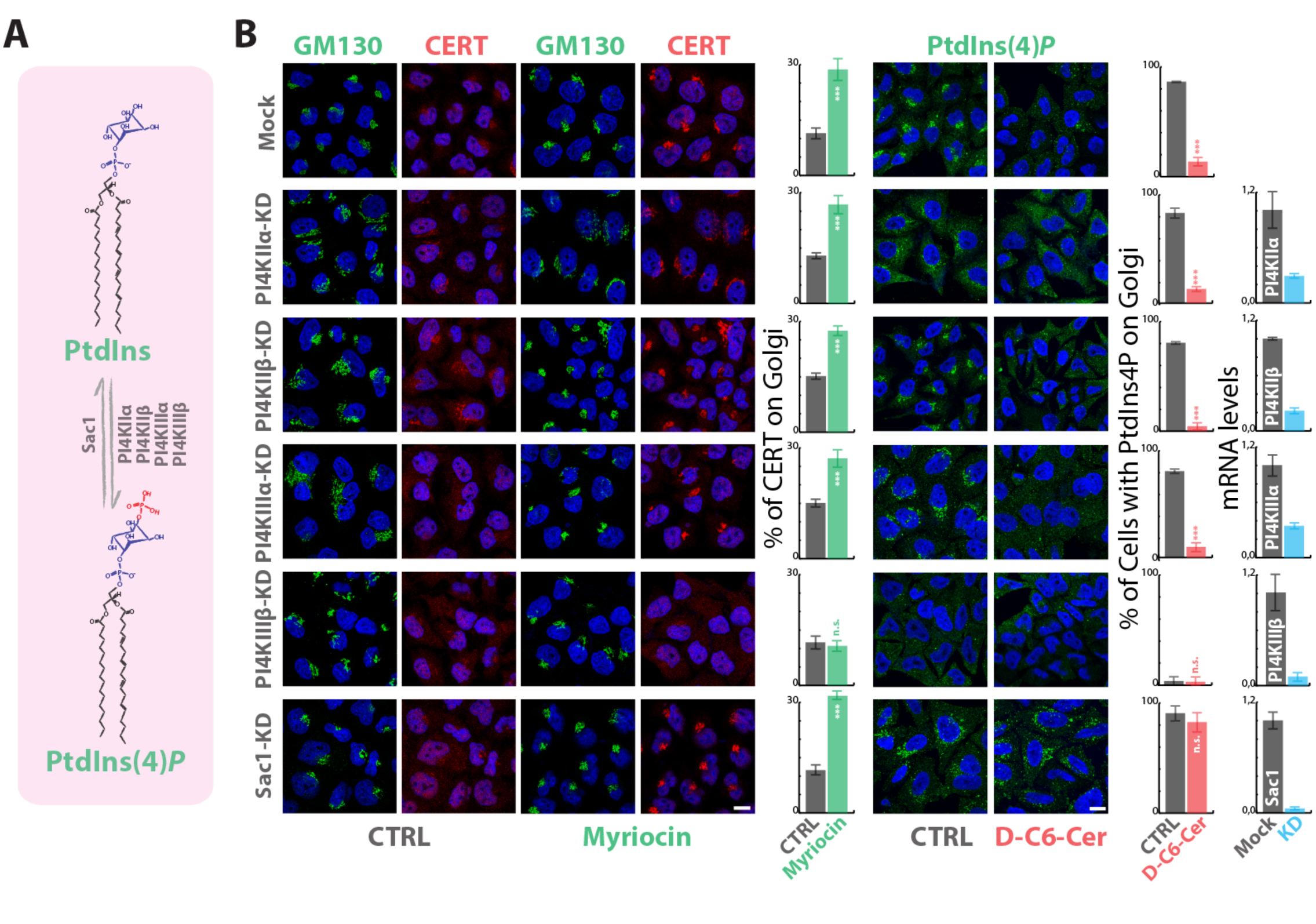
PI4KIIIβ and Sac1 control the Golgi PtdIns4P pool under SL flow. **A)** Schematic representation of PtdIns phosphorylation by PI4Ks and of PtdIns(4)*P* dephosphorylation by Sac1. **B)** Cells silenced for the expression of PI4KIIα, PI4KIIβ, PI4KIIIα, PI4KIIIβ, or SAC1 were treated with myriocin (2,5 μM; 24 hours; left panels) or with D-C6-Cer (10μM; 60 min; right panels). CERT recruitment to the Golgi region was evaluated by immunofluorescence and quantitated as in **Figure 1C**. PtdIns(4)*P* consumption under D-C6-Cer (10μM, 1 hour) was evaluated as in **Figure 4A**. The KD efficiency for PI4Ks and SAC1 is assessed by RT qPCR (right histograms). Bars, 10 μm. Data are means ± S.E.M, of at least 3 independent experiments.

PI4KIIIβ can be phosphorylated by PKD on Serine 294 and this phosphorylation stimulates 4-kinase activity (Hausser, Storz et al., 2005). We thus wondered whether PI4KIIIβ is phosphorylated by PKD in response to SL flow. As shown in **Fig EV12A** C6-D-Cer treatment induced a fast and transient PI4KIIIβ phosphorylation (as recognized by the anti-PKD pMOTIF antibody), which is completely prevented by treatment with the PKD inhibitor Gö 6976. Moreover, when the PI4KIIIβ-S294A mutant was tested for PKD-dependent phosphorylation following C6-D-Cer treatment it was found to be non-phosphorylated suggesting that Ser294 is the only phosphosite in PI4KIIIβ to be phosphorylated by PKD in response to sustained SL flow (**Fig EV12B**). Nevertheless, while PI4KIIIβ phosphorylation by PKD on Ser294 leads to enzyme activation and increased PtdIns(4)*P* production (Hausser et al., 2005) (**Fig EV12C**), the net effect of PKD activation by SLs is, in fact, a decrease in TGN PtdIns(4)*P* levels. We thus considered the involvement of Sac1 phosphatase.

Sac1 is a transmembrane protein cycling between the ER and the Golgi complex (Blagoveshchenskaya et al., 2008). It has been shown that signalling cues influence the distribution of Sac1 between these two compartments (Bajaj Pahuja et al., 2015) with serum starvation inducing its translocation to the Golgi complex and consequent TGN PtdIns(4)*P* consumption (Blagoveshchenskaya et al., 2008). We asked whether increased SL flow would mimic this situation. To test this hypothesis, cells expressing a HA-tagged version of Sac1 were treated with C6-D-Cer, and changes in Sac1 subcellular distribution associated with PtdIns(4)*P* consumption were evaluated. As shown in **Fig EV13A**, Sac1 overexpression sensitizes cells to C6-D-Cer treatment, since low C6-D-Cer concentrations (1μM) caused the disappearance of PtdIns(4)*P* at the TGN. Nevertheless Sac1 intracellular distribution was not influenced by the treatment with most of the protein being ER localized even after C6-D-Cer administration. Moreover, Sac1 overexpression was sufficient to counteract CERT recruitment to the Golgi complex induced by myriocin treatment (**Fig EV13B**). Importantly, Sac1-KD rendered TGN PtdIns(4)*P* resistant to C6-D-Cer treatment indicating that C6-D-Cer induced PtdIns(4)*P* decrease results from its Sac1-operated conversion to PtdIns (**Fig 6B**).

Thus we concluded that the metabolic couple PI4KIIIβ/ Sac1 is responsible for the control of PtdIns(4)*P* levels at the TGN induced in response to SL flow and consequent PKD signalling and that this control does not require Sac1 relocation to the Golgi complex.

### SL flow induces PKD-dependent OSBP1 phosphorylation and OSBP1-dependent PtdIns(4)P consumption

Sac1 is not known to be a PKD substrate and our attempts to detect PKD dependent Sac1 phosphorylation gave negative results, thus we considered the possibility that PKD could phosphorylate an intermediate factor that would then present PtdIns(4)*P* to Sac1. Recent publications have indicated that the PtdIns(4)*P* effector, OSBP1 interacts with Sac1 (Wakana, Kotake et al., 2015), it is phosphorylated by PKD (Nhek, Ngo et al., 2010), and is able to transfer PtdIns(4)*P* from the TGN to the ER channelling this lipid for Sac1-dependent dephosphorylation (Mesmin et al., 2013). We, thus, decided to investigate the possible role of OSBP1 in the SL-induced PtdIns(4)*P* consumption.

First, we tested whether SL flow induces OSBP1 phosphorylation by PKD. To this aim, cells expressing OSBP1-GFP were treated with C6-D-Cer, and PKD phosphorylation was revealed by pMOTIF antibody after immunoprecipitation. As shown in **Fig 7A** PKD-mediated OSBP phosphorylation started increasing after 15 min of C6-D-Cer administration and remained high at 60 min treatment while pre-treatment of cells with the PKD inhibitor Gö 6976 completely abolished these dynamics. Noteworthy, OSBP is phosphorylated by PKD in response to SL flow with a delayed and persistent dynamic when compared with PI4KIIIβ (compare with **Fig EV12A**) or with general PKD substrates (compare with **Fig 4C**). While the molecular reason for this difference remains unclear the fact that OSBP1 phopshorylation is maximal at time points when PtdIns(4)*P* is consumed suggested that OSBP1 mediates PtdIns(4)*P* consumption in response to C6-D-Cer administration.

**Figure 7:**
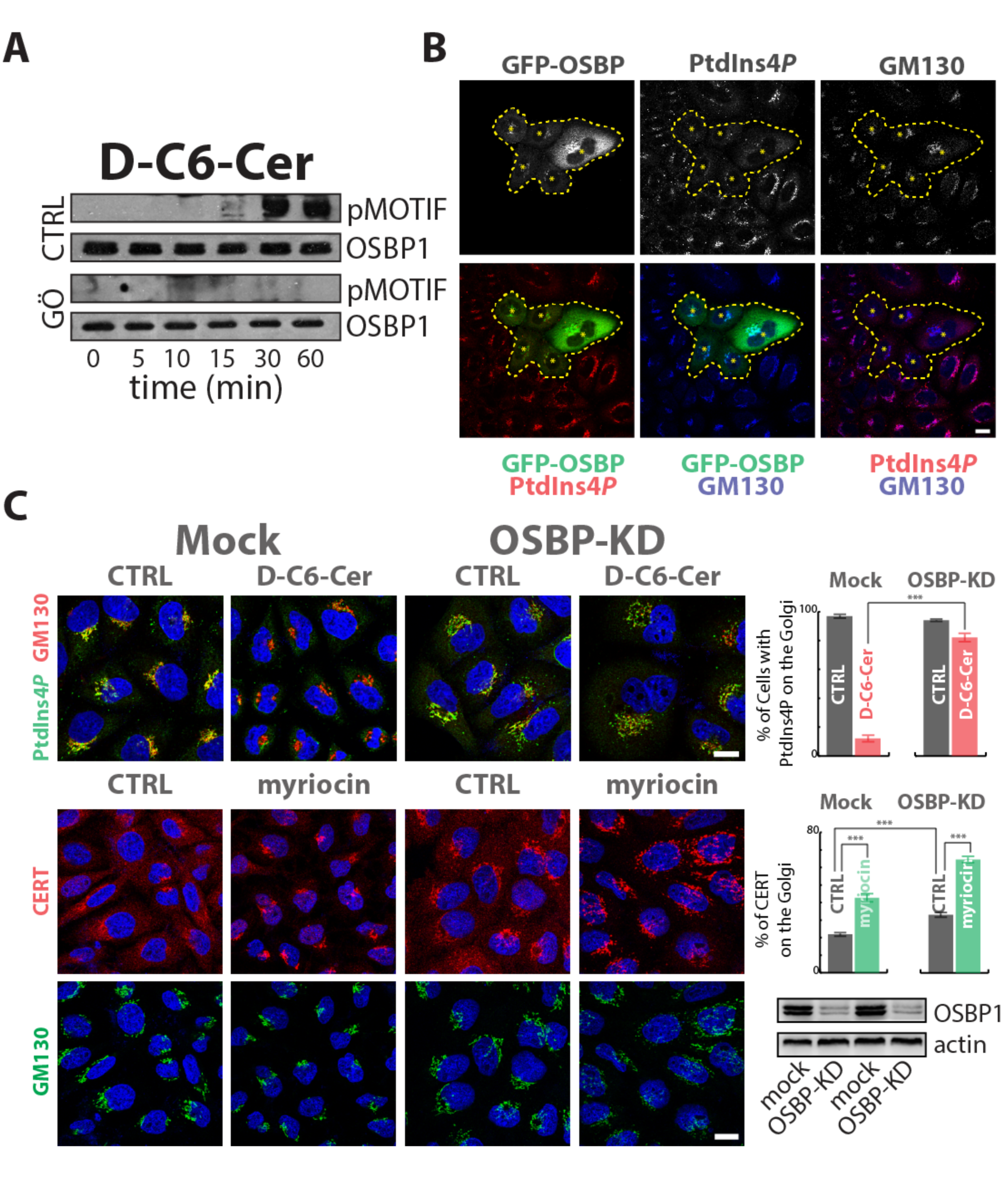
OSBP1 mediates SL flow induced PtdIns(4)P consumption. **A)** Cells expressing OSBP1-GFP were treated for the indicated times with D-C6-Cer (10μM) lysed, and protein lysates were subjected to immunoprecipitation by the use of anti-GFP antibody, SDS-PAGE and Western Blotting. PKD dependent phosphorylation of OSBP was revealed by pMOTIF antibody. The PKD inhibitor Gö 6976 (10 μM, pre treatment 1 hour) was used as a specificity control. **B)** Cells transiently expressing OSBPGFP were fixed and permeabilized as in **Figure 3** and stained with anti-GM130 (blue), and anti-PtdIns(4)*P* (red). Asterisks and dashed line indicate OSBP-GFP transfected cells. **C)** Cells KD for OSBP1 were treated with D-C6-Cer (10μM) for 1 hour, fixed, permeabilized, and stained as in **Figure 3A** (upper left panels). The percentage of cells showing PtdIns(4)*P* at the Golgi in the different conditions tested is reported (upper right panel). Cells KD for OSBP1 were treated with myriocin (2,5 μM for 24 hours) fixed, permeabilized and stained for CERT (red) and GM130 (green). CERT recruitment to the Golgi region was evaluated by immunofluorescence and quantitated as in **Figure 1C** (middle right panel). OSBP1 KD efficiency was evaluated by Western Blotting (lower right panel). Bars, 10 μm. Data are mean ± standard deviations of at least three independent experiments.

We thus tested if OSBP1 has a role in PtdIns(4)*P* consumption induced by sustained SL flow. First we confirmed that OSBP1 overexpression was sufficient to induce the consumption of PtdIns(4)*P* at the TGN (Mesmin et al., 2013) (**Fig 7B**). Importantly when OSBP1-KD cells were subjected to C6-D-Cer treatment PtdIns(4)*P* was protected from SL flow induced consumption (**Fig 7C**). Moreover OSBP1-KD, similar to what observed in SMS1-KD, induced CERT recruitment to the TGN under basal conditions (**Fig 7D**).

Taken together these data are compatible with a model whereby the SL-induced PKD-dependent phosphorylation of OSBP1 stimulates the non-vesicular transport of PtdIns(4)*P* to the ER for its dephosphorylation operated by Sac1.

## Discussion

The *post*-Golgi membranes enrichment in SLs and sterols is a shared and fundamental property of eukaryotic cells and has a key role in shaping both cell response to the environment (by influencing cell signalling) and intracellular organization (by determining intracellular protein distribution) (Holthuis & Menon, 2014, Lingwood & Simons, 2010, Patterson et al., 2008, van Meer et al., 2008). When considering the establishment of the *post*-Golgi membrane territory two traffic fluxes need to be taken into account. The first consists of material traversing the Golgi stack in a *cis*-to-*trans* direction, mediates the transport of proteins and of the bulk of membrane lipids, and requires vesicles (Nakano & Luini, 2010). The second consists of lipids exchanged at ER-TGN MCSs (De Matteis & Rega, 2015, Holthuis & Menon, 2014) where lipid-transfer-proteins execute the non-vesicular transport of selected lipid species (including SLs, and sterols). These two fluxes are orthogonal as they are operated by distinct machineries, still they converge at the TGN and need to be coordinated in order for TGN derived carriers to transport the proper protein and lipid cargo to *post*-Golgi destinations.

A fundamental factor regulating both non-vesicular lipid transport and formation of TGN derived carriers is PtdIns(4)*P* (D‘Angelo et al., 2008). PtdIns(4)*P* indeed, is the recruitment factor used by the lipid transfer proteins CERT, FAPP2, and OSBP1 to deliver SLs and cholesterol to TGN membranes (D‘Angelo et al., 2008). PtdIns(4)*P* also recruits to TGN cytosolic factors able deform membranes (i.e., FAPP1, Arfaptins, AP1/ Clathrin) thus promoting membrane budding (Cruz-Garcia, Ortega-Bellido et al., 2013, Godi et al., 2004, He, Scott et al., 2011, Lenoir, Coskun et al., 2010, Wang, Wang et al., 2003), and mediates the association of nascent carriers with actin cytoskeleton, which provides the tensile force required for their formation (i.e., via GOLPH3 recruitment) (Dippold, Ng et al., 2009). As a combination of these activities PtdIns(4)*P* controls both the formation of TGN derived membrane carriers and their lipid composition.

In addition lipid remodelling and carriers formation at the TGN are both coordinated by signalling mechanisms that converge on PKDs (Malhotra & Campelo, 2011). PKDs, indeed, are recruited to TGN by ARF1 and DAG (Malhotra & Campelo, 2011) to phosphorylate and activate PI4KIIIβ (Hausser et al., 2005) allowing its interaction with 14-3-3γ and CtBP1/BARS in a complex required for carriers fission (Pagliuso, Valente et al., 2016, Valente, Turacchio et al., 2012). At the same time Cer transported to the TGN by CERT, is converted to SM in a reaction that generates DAG. TGN-produced DAG has, in turn, the potential to activate PKDs thus establishing a positive feedback in SM synthesis and *post-*Golgi carriers formation (Malhotra & Campelo, 2011, Thomaseth, Weber et al., 2013, Villani, Subathra et al., 2008). A further PKD substrate is CERT itself, where its phosphorylation reduces its affinity for PtdIns(4)*P* (Fugmann, Hausser et al., 2007) in a process that promotes CERT cycling and results is a further positive feedback loop between CERT transfer activity and PKD activity (Weber, Hornjik et al., 2015).

According to this description the TGN is equipped with a basal lipid remodelling system accounting for the enrichment of *post-*Golgi carriers (and thus *post-*Golgi compartments) in SLs and cholesterol (Deng et al., 2016). But how does this system react to acute fluctuations in SL production/ flow? While, indeed, both reduced and excessive SM synthesis at the TGN have been reported to impair TGN to plasma trafficking (Duran et al., 2012, Subathra, Qureshi et al., 2011), a comprehensive picture on how SL metabolic oscillations are matched to TGN and *post-*Golgi SL composition was missing. Here we describe a molecular circuit linking SL flow to the regulation of PtdIns(4)*P* turnover at the TGN.

We indeed show that **(i)** SL flow stimulates the consumption of the TGN localized PtdIns(4)*P*; **(ii)** as a consequence, SL flow negatively regulates CERT, FAPP2, and OSBP1 recruitment to the Golgi complex (and thus Cer, GlcCer, and cholesterol transport to the TGN); **(iii)** which results in SL flow repressing the production of SM and GSLs at the TGN. Specifically we show that a sudden increase in SM synthesis at first induces the PKD-dependent phosphorylation of PI4KIIIβ with a possible transient increase in PtdIns(4)*P* production. Subsequently PKD phosphorylates OSBP1 inducing PtdIns(4)*P* relocation to the ER and Sac1 dependent PtdIns(4)*P* dephosphorylation (**Fig EV14A**). Given the role of PtdIns(4)*P* in promoting SM and GSL synthesis at the TGN, sustained SM synthesis inhibits SM and GSL production at the TGN (**Fig EV14B**). Importantly this circuit appears to be active also under steady-state conditions as reductions in SL flow or SM synthesis translate in reduced basal PKD activity and recruitment of PtdIns(4)*P* effectors to the TGN.

Based on these data we envisage a model where acute fluctuations in SL provision are buffered by a circuit that acts as a homeostatic device to keep SM and GSLs synthetic rates constant irrespective of instant changes in the SL flow. In other words, the circuit described here would work in a way similar to a voltage stabilizer in a power supply, which generates a fixed output voltage of a pre-set magnitude regardless of changes to its input. The intuitive outcome of the operation of this circuit would be that of providing robustness and stability to the TGN lipid biosynthetic system and, as a consequence, that of maintaining constant the SM, GSL and possibly cholesterol composition of TGN and *post-*Golgi compartments. Interestingly, when considered in the general frame of cellular lipid homeostatic systems, the circuit described here presents some unique features: **(i)** it relies on the local sensing of a transient event (i.e., SM synthetic rate at the TGN) and not on the detection of global mass levels of a given metabolite (Holthuis & Menon, 2014); **(ii)** the architecture of this circuit wires together the local metabolism of lipid mediators with roles in metabolic and trafficking TGN functions (namely Cer, SM, DAG, PtdIns(4)*P*, GSLs, and cholesterol) (Holthuis & Menon, 2014) (De Matteis & Luini, 2008). It is thus tempting to speculate that by these unique features this circuit preforms a harmonizing function on TGN lipid remodelling during the dynamic event of *post-*Golgi carriers formation.

In conclusion, decades of research in membrane biology have shown how membrane composition impacts on membrane properties (Lingwood & Simons, 2010, van Meer et al., 2008). Specifically SLs and cholesterol at the TGN and *post*-Golgi membranes exert an organizing function on other lipids and membrane proteins influencing their lateral distribution, their trafficking, and ultimately their specific functions (Holthuis & Menon, 2014). Previous studies have described homeostatic circuits devoted to maintain the global SL content in eukaryotic cells by regulating the reactions initiating SL production in the ER (Breslow, Collins et al., 2010, Cantalupo et al., 2015, Vacaru et al., 2009). In this study we provide evidence for the existence (and dissect the architecture) of a homeostatic circuit accounting for the localized and acute control of SL metabolism at the TGN and thus of *post*-Golgi membranes composition.

## Acknowledgments

We thank MA. De Matteis for initial discussions, C. Luberto, D. Corda, and A. Colanzi for critically reading the manuscript. We thank the IBP-bioimaging facility for technical support. GD’A acknowledges the financial support of AIRC (MFAG 10585), of the Italian Ministry of Health (GR-2011-02352256) and of MIUR (PON_00862). RR acknowledges the financial support of FIRC (fellowship N° 15111). AL acknowledges FFC [Italian Cystic Fibrosis Research Foundation FFC #2 2014], MIUR [COSM Progetto Interomics], AIRC [Italian Association for Cancer Research, IG 15767] the MIUR Project “FaReBio di Qualità”, the PON projects no. 01/00117 and 01-00862, PNR-CNR Aging Program 2012–2014, and Progetto Bandiera ‘Epigen’. VM acknowledges support of the Ministerio de Economía y Competitividad, “Centro de Excelencia Severo Ochoa 2013-2017,” SEV-2012-0208. VM is an Institució Catalana de Recerca i Estudis Avançats professor at the Center for Genomic Regulation, and the work in his laboratory is funded by grants from Ministerio de Economía y Competitividad’s Plan Nacional (ref. BFU2013-44188-P).

## Contributions

GD’A supervised the entire project, with advice from AL and PS; GD’A and SC wrote the manuscript with comments from all co-authors; SC, with the help of LS, RR, MP, and DR, designed and conducted all the experiments described. ND designed the strategy and produced plasmid vectors for SMS1 and GCS. SC designed the strategy and produced plasmid vectors for Sac1. FC, and JvG performed experiments described in **Fig EV5, EV6**, **and EV11A** under the supervision of VM. AH provided the PI4KIIIβ-S294A plasmid used in **Fig EV12B**.

## Competing financial interests

The authors declare no competing financial interests.

## Methods

## Reagents

N-C6:0-D-*erythro*-Sphingosine (D-C6-Ceramide), N-Hexanoyl-L-*erythro*-sphingosine (LC6-Ceramide), D-*erythro*-2-N-[6‘-(1"-pyridinium)-hexanoyl]-sphingosine bromide (LCL29) were from Matreya, USA. D-*erythro*-Sphingosine (D-Sph) was from Avanti Polar, USA. Phorbol 12,13-dibutyrate (PdBU) and GÖ 6976 were from Sigma-Aldrich. PIK93 inhibitor was from Echelon Bioscience, USA. Fumonisin B1 (FB1) and myriocin were from Enzo Life Sciences, USA. Digitonin was from Calbiochem, USA. The anti-CERT, anti-OSBP polyclonal antibodies and the anti-rabbit and anti-mouse IgG Cy3-conjugated antibodies were from Sigma-Aldrich, USA. The anti-HA.11 clone 16B monoclonal antibody was from Covance, USA. The anti-GOLPH3 polyclonal and Mouse anti-GFP antibodies were from Abcam, UK. The anti-GM130 polyclonal antibody was from NOVABIOS, IT. The anti-PtdIns(4)*P* mouse monoclonal IgM antibody was from Echelon Bioscience, USA. The anti-Golgin-97, anti- FAPP2, anti-GST polyclonal antibodies were a kind gift of Antonella De Matteis laboratory. The sheep anti human TGN46 was from Bio-Rad, USA, and monoclonal mouse anti γ-adaptin (clone 100/3) was from Sigma-Aldrich, USA. The anti-PKD, anti-pPKD (Ser916), and anti-(Ser/Thr) PKD substrates (pMOTIF) polyclonal antibody were from Cell Signaling Technology, USA. Donkey anti-rabbit IgG Alexa Fluor-488 conjugated, Donkey anti-rabbit IgG Alexa Fluor-568 conjugated Donkey anti-mouse IgM Alexa Fluor-488 conjugated, Donkey anti-mouse IgM Alexa Fluor-568 conjugated for immunofluorescence microscopy were from Thermo Fisher Scientific, USA. All other unmentioned reagents were from Sigma-Aldrich, USA.

## Cell lines and culture conditions

HeLa cells were obtained from the American Tissue Type Collection (ATTC, USA). HeLa cells were grown in RPMI-1640 medium (Gibco, USA) supplemented with 10% (v/v) foetal calf serum (FCS) (Gibco), containing 4.5 g/L glucose, 2 mM L-glutamine, 100U/ml penicillin and streptomycin. Cells were grown under controlled atmosphere (5% CO2 and 95% air) at 37°C.

## Construct and Plasmids

To generate the human Sac1 (NM_014016), SMS1 (NM_147156), and GCS (NM_003358) expressing plasmids total mRNA was purified, from HeLa cells, retrotranscribed and cDNAs were amplified by PCR using the following primers:

Sac1

Forward: 5 ‘- ATAGGATTCATTGAGAGAGAAGGAAGGAGGTG -3’

Reverse: 5’- ATAGATATCATTAAAAGTATGCCTGCTAATAGTG -3’

SMS1

Forward: 5’ – GCGCGCGAATTCATGAAGGAAGTGGTTTATTGGTC -3’

Reverse: 5’ – TGTGTCATTCACCAGCCGGCT -3’

GCS

Forward: 5’ – GCGCGCGAATTCATGGCGCTGCTGGACCTGG -3’

Reverse: 5’ – TACATCTAGGATTTCCTCTGC -3’

PCR reactions were performed as follows: 1 min at 98°C, followed by 30 cycles at 98 °C for 20 s, 60 °C for 30 s, 72 °C for 2 min 120 s and a final extension at 72 °C for 7 min. The PCR generated fragment for Sac1 was digested with BamHI and EcoRV restriction enzymes (New England Biolabs, USA) to be cloned into pcDNA4b-3xHA-expression vector (Clontech, USA). The PCR generated fragment for SMS1 and GCS were digested with EcoRI and XhoI (New England Biolabs, USA) and cloned in pcDNA4b-3xHA-expression vector (Clontech, USA). All the obtained plasmids were sequenced before use.PKD-wild type, PKD-C1α and PKD-KinDead were provided by Vivek Malhotra (Centre Genomic Regulation-CRG, Barcellona); GOLPH3-GFP was provided by Alberto Luini (Istitute of Protein Biochemistry, National Research Council, Naples, Italy). FAPP2-PH-GFP and diFAPP-PH-R18L-GFP were from Antonella De Matteis laboratory (Telethon Institute for Genetics and Medicine), ORP5-PH-GFP, and ORP8L-PH-GFP were from Pietro De Camilli laboratory (Yale Medical School); PLCδ-PH-GFP was from Roman Polishchuk laboratory (Telethon Institute for Genetics and Medicine). PI4KIIIβ-GFP, FYVE-GFP, CERT-GFP, and OSBP-GFP expressing plasmids were obtained from Daniela Corda laboratory (Institute of Protein Biochemistry, National Research Council, Naples, Italy) PI4KIIIβ-S294A-GFP was obtained from Angelika Hausser laboratory (University of Stuttgart, Institute of Cell Biology and Immunology, Germany).

### siRNA treatments and transfections

The siRNAs for human SMS1 (NM_147156.3), CERT (NM_001130105.1), GCS (NM_003358), PKD1 (NM_002742.2), PKD2 (NM_001079880.1), PKD3 (NM_005813.4), OSBP1 (NM_002556.2), PI4KIIα (NM_018425), PI4KIIIα (NM_058004), PI4KIIβ (NM_018323), PI4KIIIβ (NM_001198773), Sac1 (NM_014016), (**Table S1**) were obtained from Sigma-Aldrich (Italy). HeLa cells were plated at 20% confluence in 24-well plates on 24-mm coverslips, or in 12-well plates, and transfected with 120 舁pmol of siRNAs with Oligofectamine (Invitrogen, USA), according to the manufacturer protocol. 72舁hours after the initial treatment with the siRNAs, the cells were processed for the different experiments. The silencing efficiencies were evaluated either by Western Blotting or by qPCR using specific primers (**Table S2**).

The constructs SacI-HA, PKD-wt-GST, PKD-KinDead-GST, PKD-C1α-GST, OSBP-GFP were transfected into HeLa cells using TransIT-LT1 Transfection Reagent (Mirus Bio LLC, USA), according to the manufacturer instructions.

### RNA extraction and real-time PCR

Total RNA was isolated from HeLa cells using RNeasy Mini kits (Qiagen, Germany) according to the manufacturer instructions. The yield and the integrity of the RNA were determined using a spectrophotometer (NanoDrop 2000c; Thermo Scientific, USA), by TAE agarose gel electrophoresis, and with an Agilent 2100 Bioanalyser (Agilent Technologies, USA). RNA (1 μg) was reverse transcribed using QuantiTect Reverse Transcription kits (Qiagen, Germany) according to the manufacturer instructions and subjected to real-time qPCR with gene-specific primers (**Table 2**) in the presence of LightCycler^®^ 480 SYBR Green I Master Mix (Roche, Switzerland) on a LightCycler^®^ 480 II detection system (Roche, Switzerland). Relative abundance of mRNA was calculated by normalization to hypoxanthine phosphoribosyltransferase -1 (HPRT1).

**Table 1.**
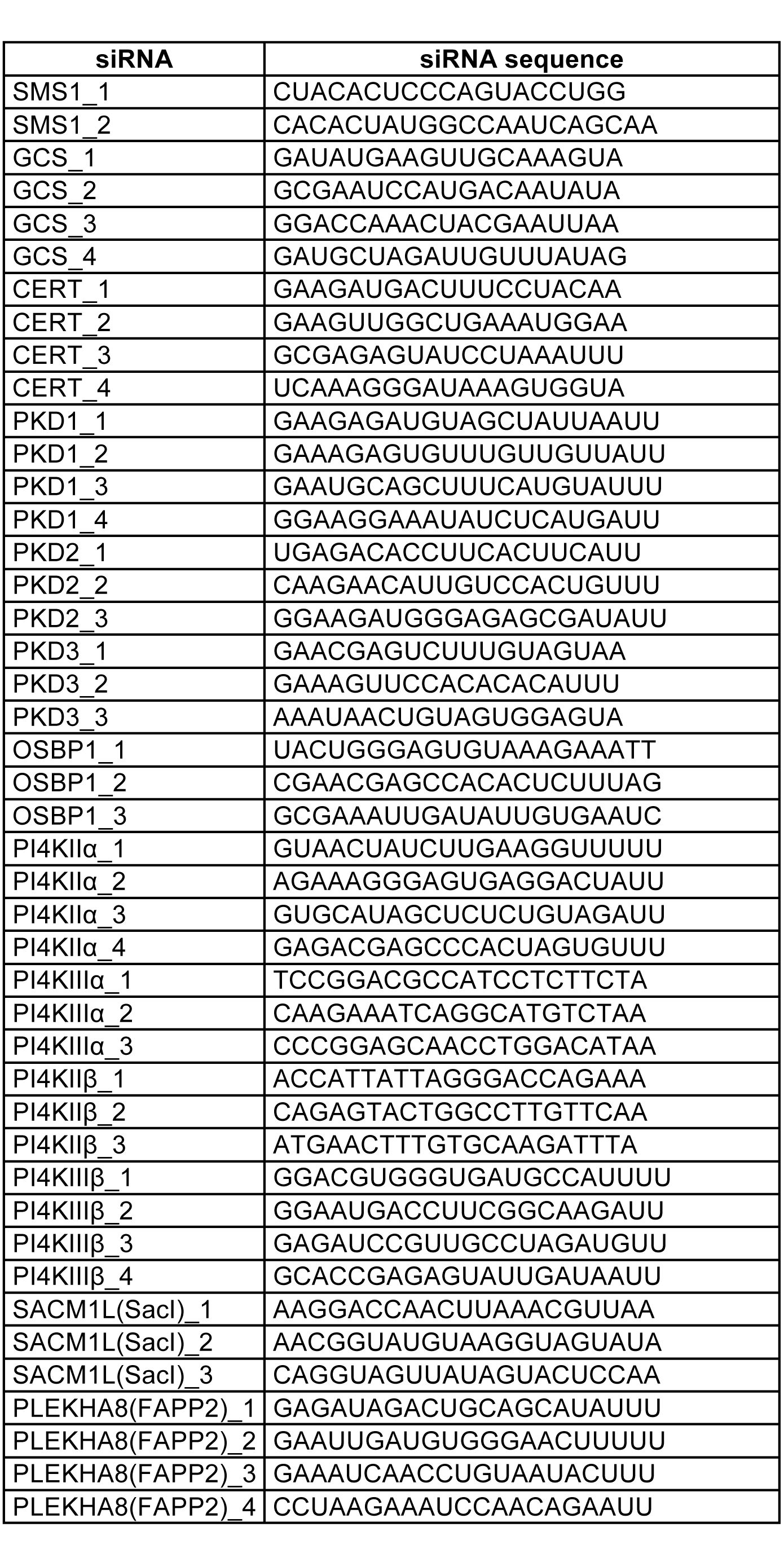
Sequences of siRNA used in this study

**Table 2.**
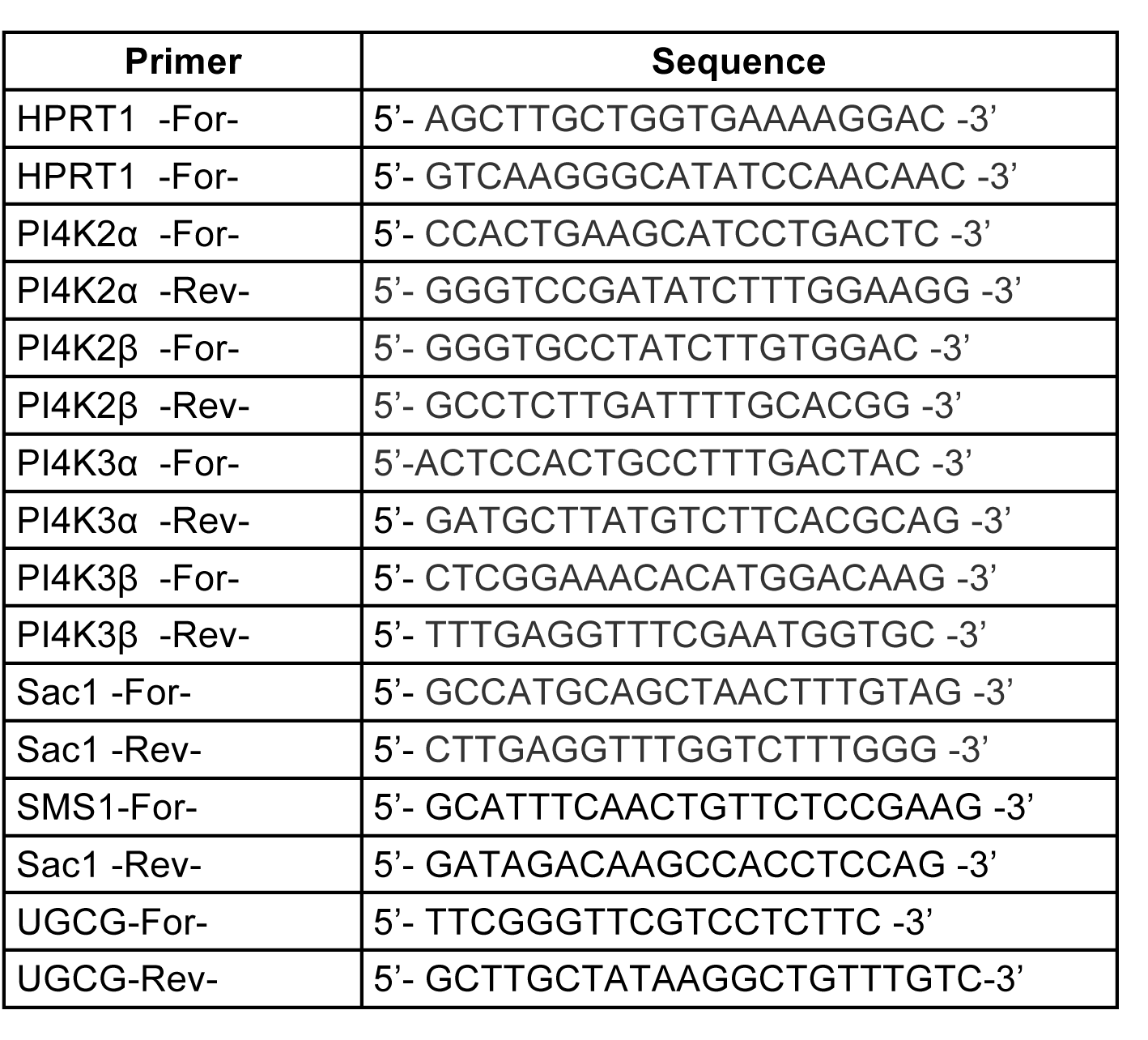
Sequences of primers used for Real Time q-PCR in this study

## Immunoblotting

Proteins were detected using the mentioned primary antibodies. The signals were visualized using a goat anti-rabbit or goat anti-mouse IgG-HRP secondary antibody (Santa Cruz, USA) at a 1:10,000 dilution.

## Immunoprecipitation assays

HeLa cells transfected with the indicated GFP-tagged plasmids were lysed in a buffer consisting of: 20 mM MOPS pH 7,0, 2 mM EGTA, 5 mM EDTA, 60 mM βglycerophosphate, 30 mM NaF, 1 mM Na3VO4, 1mM DTT, 1% (v/v) Triton-X100, Protease inhibitor. Total lysates (1mg) were mixed with 1 μg of anti-GFP IgGs O.N., and incubated for 1h with Protein A-Sepharose (Sigma). After centrifugation and washing steps proteins associated with the precipitated fraction were eluted by adding Sample Buffer (0.125 M Trizma base, 4% SDS, 20% glycerol, 10 % β-mercaptoethanol, pH 6.8), and the eluted fraction was analysed by SDS-PAGE and Western blotting.

## Sphingolipid metabolic measurements

HeLa cells pre-treated with myriocin (2,5μM) or FB1 (50μM); were pulse-labelled in RPMI-1640 10% FCS, with 0,1 μCi/mL (≈ 5nM) [^3^H]-D-erythro-Sphingosine or with [^3^H]-D-erythro-C6-Ceramide for 2 hours mixed with increasing amounts on non-radioactive D-Sph and C6-D-Cer respectively. After 2 hours cells were harvested and processed for lipid extractions. Lipids were spotted on silica-gel high performance-TLC (HPTLC) plates (Merck, Germany), and resolved with a mixture of chloroform: methanol: water (65:25:4 v/v/v). To visualize the unlabeled standards (i.e., Cer, GlcCer, and SM) the TLC plates were placed in a sealed tank saturated with iodine vapours, while the radiolabelled lipids were analysed using a RITA**^®^** TLC Analyser (Raytest, Germany), and quantified using GINA**^®^** (Raytest, Germany) software analysis. The percentage of total C.P.M. associated with Cer, GlcCer, and SM peaks for each of the three lipids is reported. For activity measurements cells labelled as described above were harvested, 1/5^th^ of total cell fraction was used for protein content determination while the rest was used for lipid extraction and analysis. For each analysed sample pmol of SM produced per μg of protein was calculated according to the following formula:

(C.P.M._SM_ /C.P.M. _0.1μCi SL_) × SL [pmol] / protein lysate [μg]
Where:

(C.P.M._SM_/C.P.M._0.1μCi SL_) × SL [pmol] / protein lysate [μg]

C.P.M._SM_ represents the counts per minute associated with the SM peak.

C.P.M. _0.1μCi SL_ represents the counts per minute associated with 0.1 μCi of ^3^H-labelled SL precursor.

SL [pmol] represents the amount of SL precursors fed to cells (pmoles).
protein lysate [μg] represents the amount of proteins obtained by lysing the sample (μg).

## Immunofluorescence, staining and image analysis

HeLa were grown on 24-mm coverslips and treated according to the experimental procedure were fixed with 4% paraformaldehyde for 15 min and washed once in PBS. Blocking Buffer (PBS 1X, 0,5% BSA, 0,05% Saponin, 50mM NH_4_Cl, 0,02% NaN_3_) was added to the cells for 20 min, followed by 1-hour incubation with the primary antibody. Subsequently coverslips were washed with PBS and incubated with secondary antibodies (1:400) diluted in blocking solution for 45 min, washed again, and mounted on glass microscope slides (Carlo Erba, Italy) with Mowiol.

The PtdIns(4)*P* staining was performed as described in (Hammond et al., 2009). HeLa cells were grown on 24-mm coverslips. At 80% confluence, the cells were quenched with PBS containing 50mM NH_4_Cl, fixed in 2% paraformaldehyde for 15 min. The cells were permeabilized for 5 min with 20μM digitonin in bufferA containing 20 mM Pipes pH 6.68, 137 mM NaCl, 2.7 mM KCl. HeLa cells were blocked in buffer A supplemented with 5% (v/v) FBS for 45 minutes. Anti-PtdIns(4)*P* monoclonal mouse antibody dissolved in buffer A with 5% FBS was used for incubation (1:200) for 1 hour. After washing with buffer A, anti-mouse IgM secondary antibodies dissolved in buffer A with 5% FBS (1:400) were used in incubations for 45 min. Cells were, then, post-fixed for 5 min in 2% paraformaldehyde, washed three times with PBS 50mM NH_4_Cl, washed once with water and mounted.

Immunofluorescence samples were examined under confocal laser microscope (ZEISSLSM700 confocal microscope systems; Carl Zeiss, Gottingen, Germany). Optical confocal sections were taken at 1 Air Unit. Images were analysed using either ImageJ (Schneider, Rasband et al., 2012) or CellProfiler (Carpenter, Jones et al., 2006) software.

## Live cell imaging and FRAP assays

HeLa cells plated on 35-mm glass bottom microwell dishes were from MatTech (USA), were transfected with plasmids encoding GFP-ARF1, FAPP2-PH-GFP and diFAPP-PHR18L-GFP (1ug) using TransIT-LT1 Transfection Reagent (Mirus Bio LLC, USA) according to manufacturer’s instruction and analysed 24 hours after transfection. For temperature control during live observation, the microscope was equipped with a temperature-controlled chamber, and the cells were imaged on a 37 °C stage in DMEM-HEPES-buffered pH 7.4 media using a Zeiss LSM 700 confocal microscope. Time-lapse series were acquired at 15 s intervals for a total time of 60 min. Image series were exported as single .lsm files, and processed with Imagej for analysis, single frame extraction, and conversion into QuickTime movies.

Fluorescence Recovery After Photobleaching (FRAP) experiments were performed bleaching the GFP-ARF1 associated fluorescence in the Golgi area (50 bleaching iterations) and then monitoring the fluorescence intensity recovery in the bleached area. For both live imaging and FRAP experiments normalised Golgi associated fluorescence was calculated according to the formula: **nGolgi *f* (t) *=*[Golgi *f* (t)/ Golgi *f*(0) / Cell *f* (t)/ Cell *f*(0)]*100**
where: **nGolgi *f* (t)** represents the normalised Golgi associated fluorescence;

Golgi *f* (t) and **Golgi *f* (0)** represent the Golgi associated fluorescence at time (t) and 0 respectively;

And **Cell *f* (t)** and **Cell *f* (0)** represent the whole cell associated fluorescence at time (t) and 0 respectively.

## Statistics

Error bars correspond to either SD or SEM according to the different experiments and as indicated in the figure legends. Statistical evaluations report on Student’s t test *p < 0.05, **p < 0.01, and ***p < 0.001 (ns, not significant).

---

**Expanded View Figure 1:**
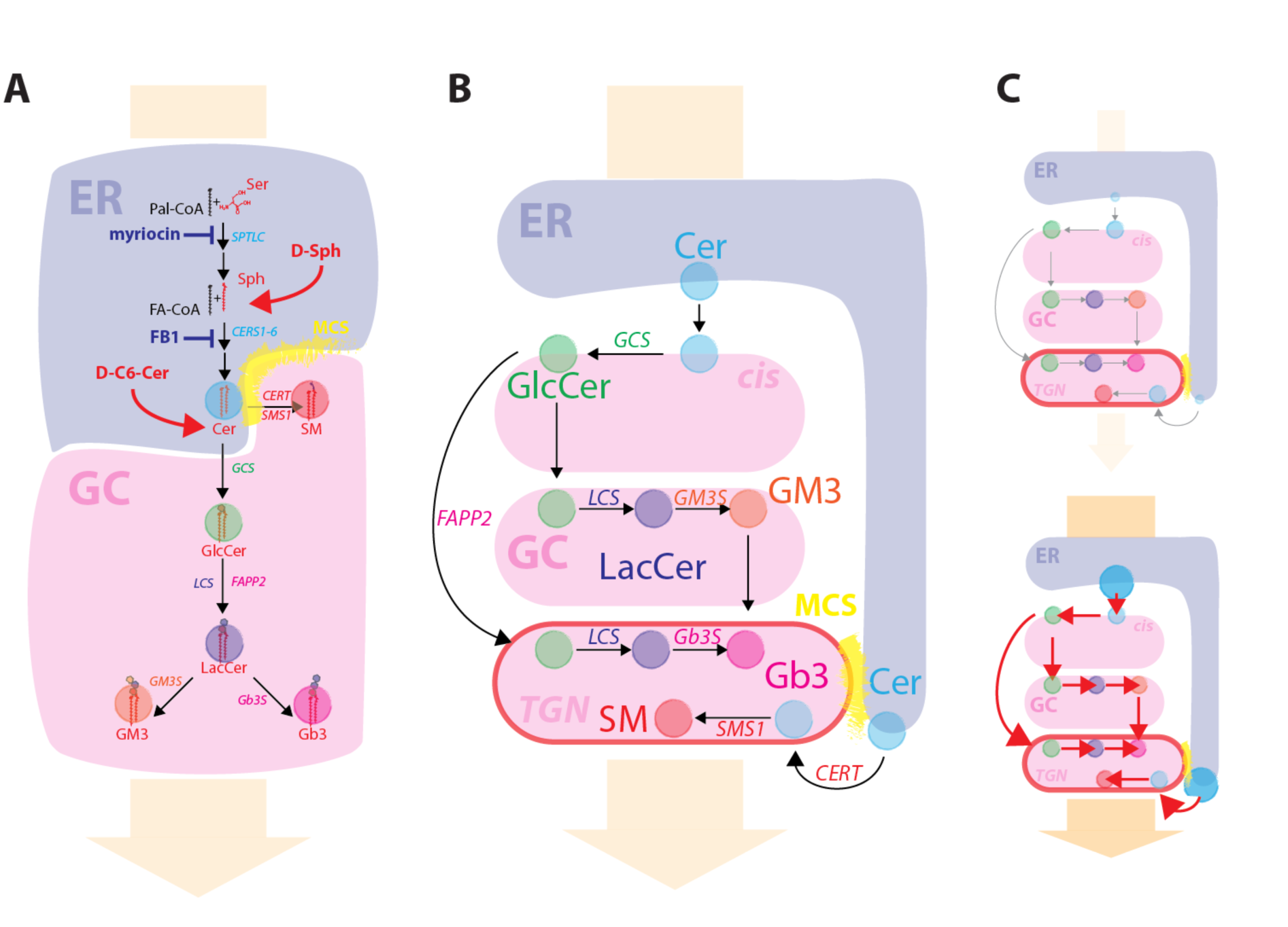
*Intracellular organization of SL metabolism* **A)** Schematic representation of SL synthetic reactions happening at the ER (blue shaded area) and at the Golgi complex (GC; red shaded area). The metabolic positions where SL synthetic inhibitors act (blue) and where cell permeable SL precursor enter SL metabolism (red) are indicated. The regions highlighted in yellow indicate ER-TGN MCSs. The orange arrow in the background indicates the direction of the biosynthetic flow. **B)** Schematic representation of the sub-Golgi distribution of SL metabolic enzymes and associated reactions. **C)** Experimental scheme for decreased (up) and increased (down) SL flow obtained by the use of cell permeable SL precursor or SL synthetic inhibitors.

**Expanded View Figure 2:**
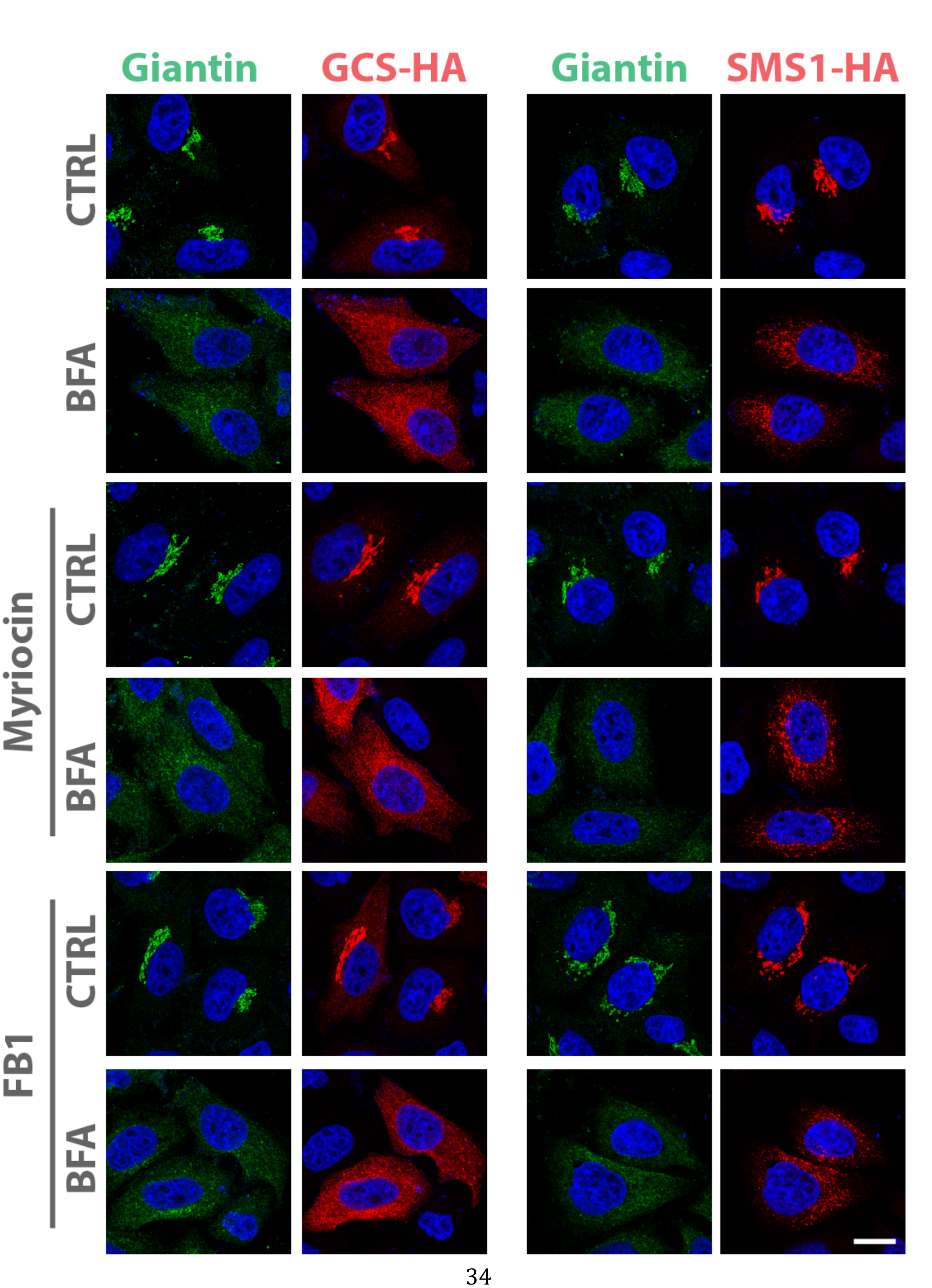
SMS1 and GCS are redistributed to the ER under BFA treatment. Cells expressing HA- tagged versions of GCS or SMS1 were either incubated with vehicle or with SL inhibitors myriocin (2.5 μM) or FB1 (50 μM) for 24 hours. Subsequently cells were treated with BFA (5μg/mL) for 30 min fixed and processed for immunofluorescence. Images show the subcellular distribution of GCS-HA and SMS1-HA (red) compared to that of the Golgi marker Giantin (green) under the different treatment conditions. Note that both GCS and SMS1 are predominantly located to the Golgi complex, and that both are redistributed to the ER under BFA administration irrespective of myriocin or FB1 treatments. Bar, 10 μm.

**Expanded View Figure 3:**
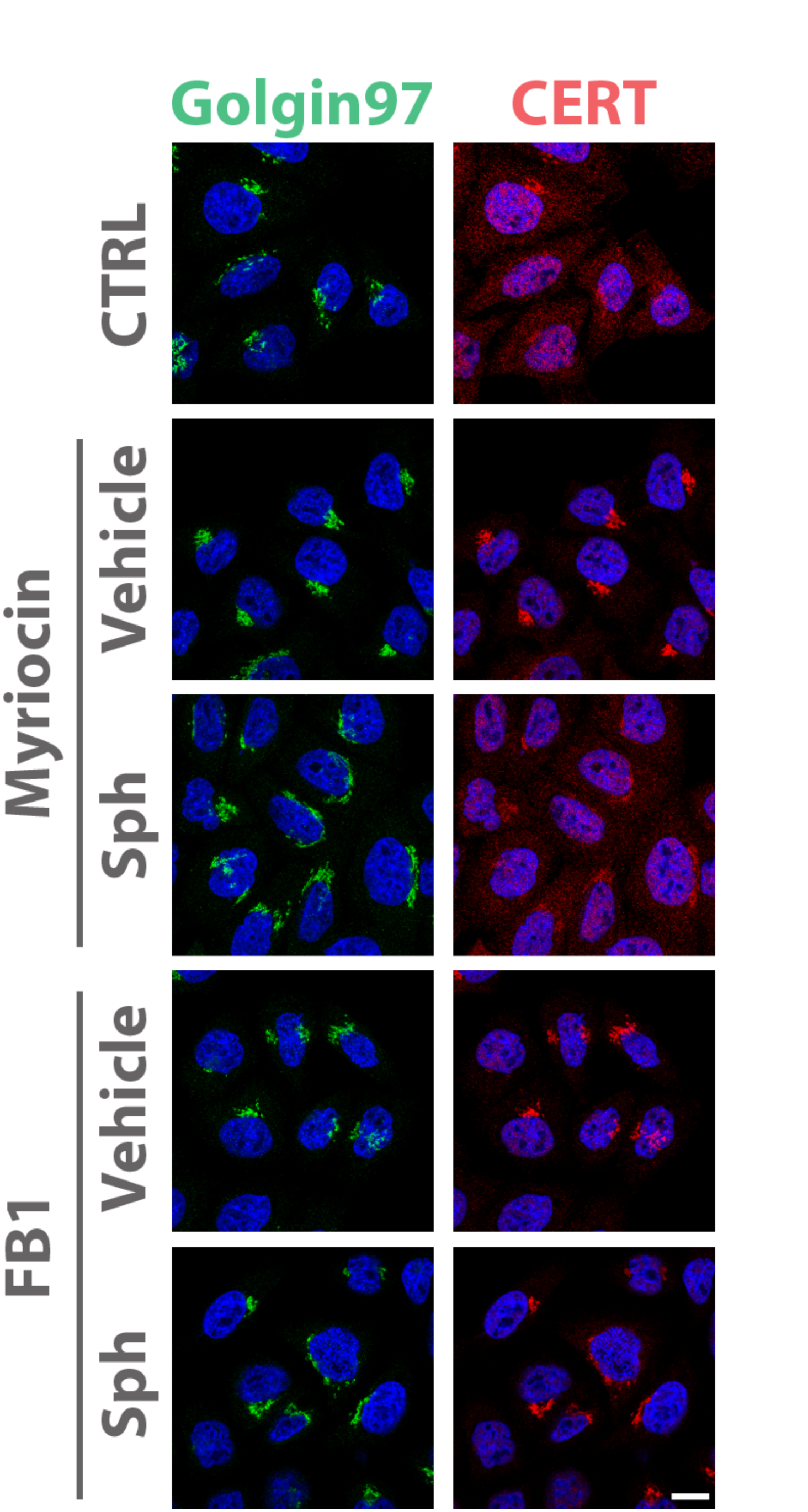
CERT association to Golgi membranes depends on SL synthetic flow. CERT localisation in HeLa cells, either non-treated (CTRL), treated with myriocin (2.5 μM for 24 hours) or FB1 (50μM for 24 hours), or treated with 30μM D-Sph for 2 hours after 24 hours treatment with myriocin or FB1. CERT is in red, the TGN marker Golgin 97 is in green, DAPI is in blue. Bar, 10 μm. Note that D-Sph treatment reverts myriocin induced CERT recruitment to the Golgi while it has no effect on FB1 induced recruitment suggesting that D-Sph conversion to D-Cer is required for CERT displacement.

**Expanded View Figure 4:**
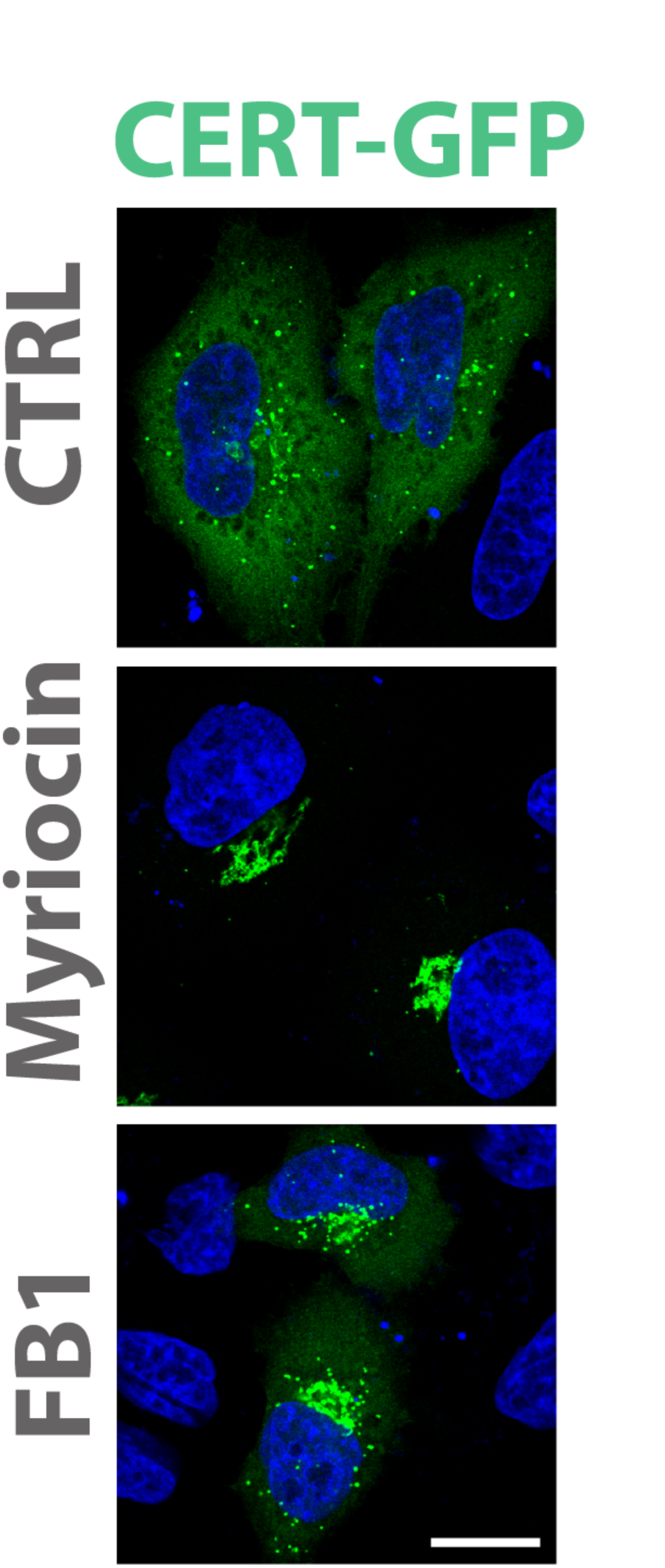
Association of GFP-CERT to Golgi membranes depends on SL synthesis. GFP-CERT localisation in HeLa cells, either non-treated (CTRL), treated with myriocin (2.5 μM, 24 hours), or FB1 (50μM, 24 hours). Bar, 10 μm.

**Expanded View Figure 5:**
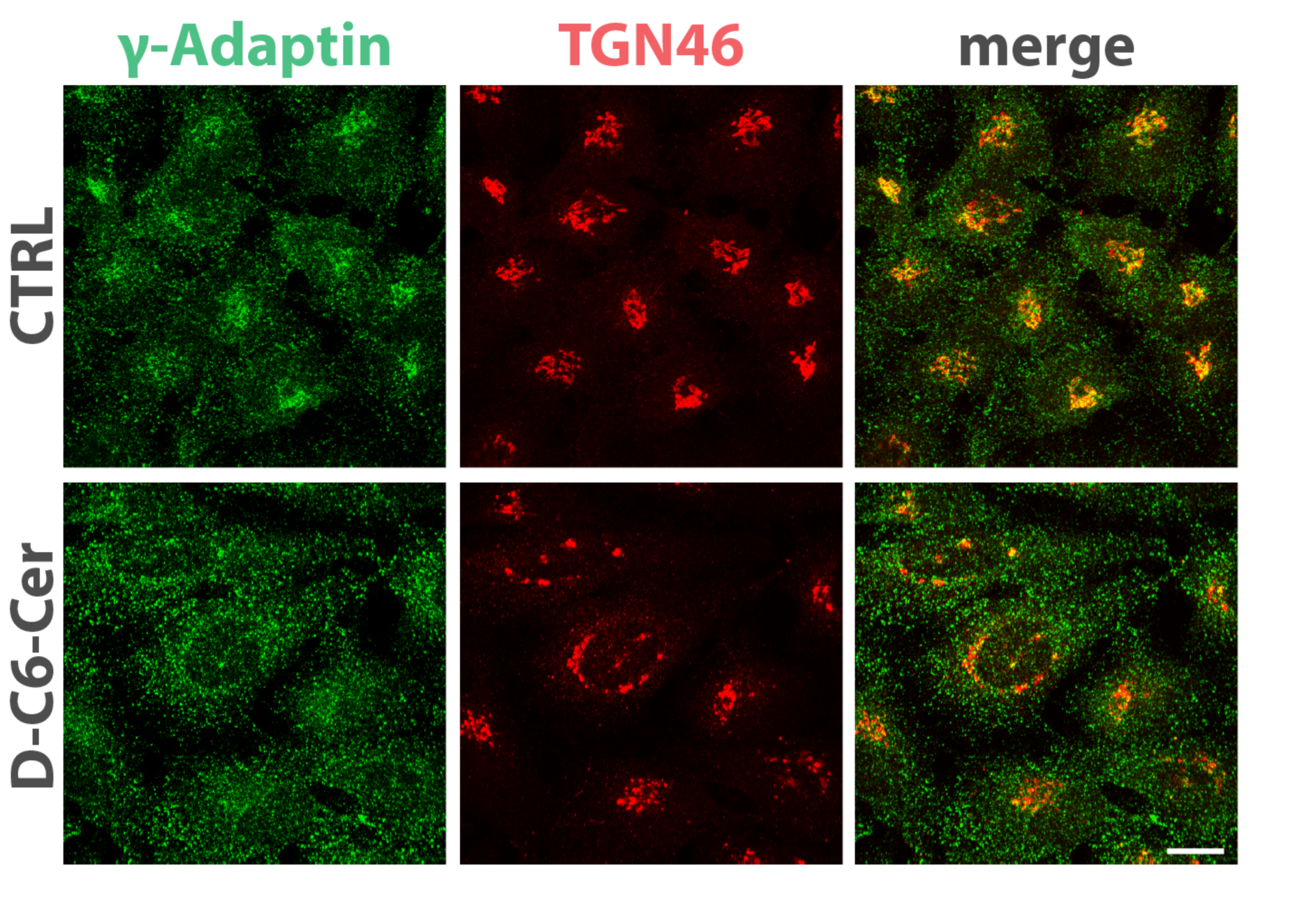
γ-Adaptin association to the Golgi is sensitive to the SL flow. Cells were treated as in **Figure 2A**, fixed and stained for γ-Adaptin (green) and TGN46 (red). Bar, 10 μm.

**Expanded View Figure 6:**
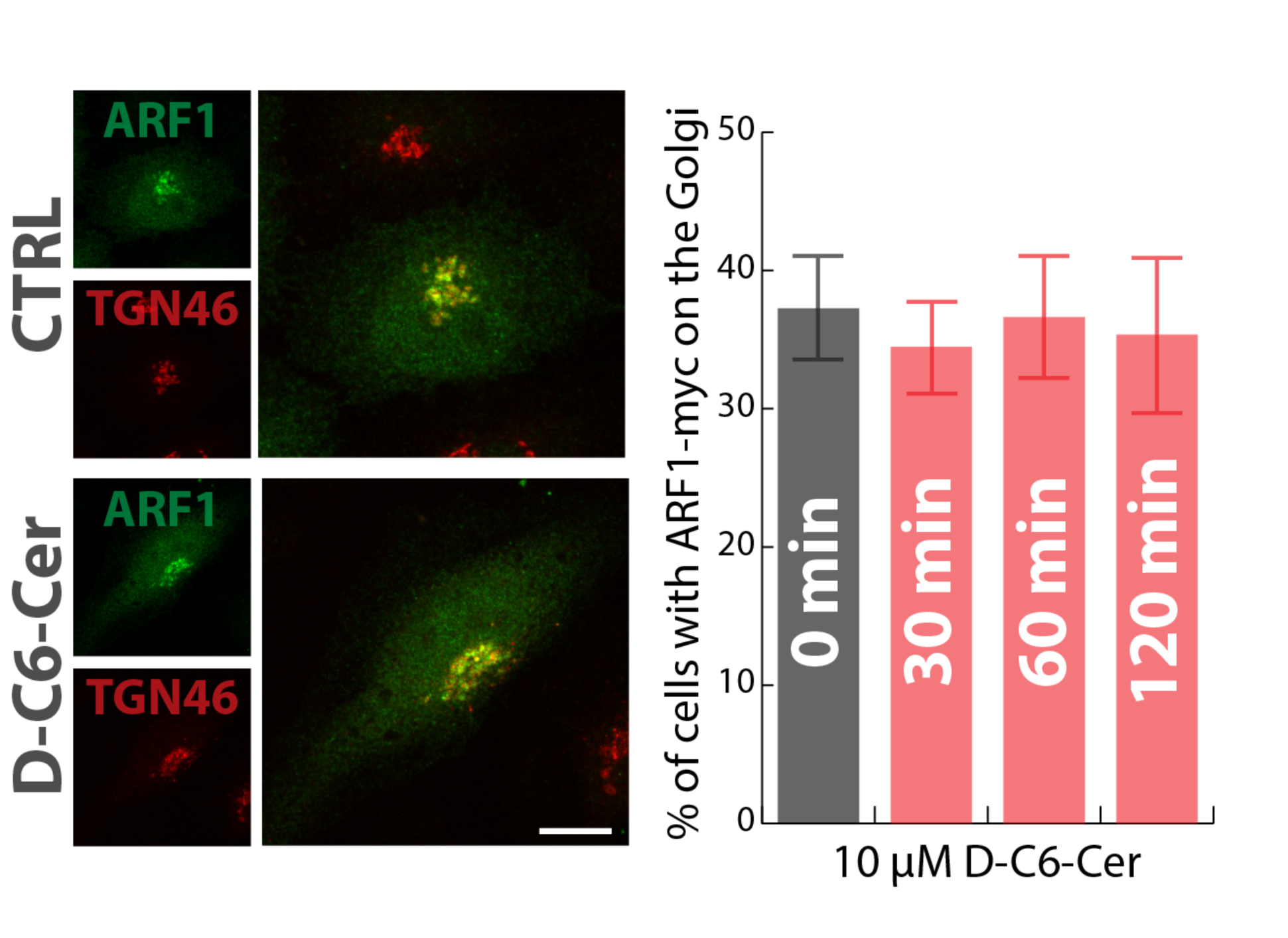
Arf1 localization to the Golgi complex is not affected by SL flow. Cells expressing a myc-tagged version of ARF1 (ARF1-myc), were treated either with EtOH or D-C6-Cer (10μM) for different times (30, 60, and 120 min). Cells were, then fixed and stained with anti-myc antibody (green) and with an anti-TGN46 antibody (red) (left panel). The percentage of transfected cells having ARF1-myc localized to the Golgi is indicated at different time points. Data are means of at least 3 independent experiments ± standard deviation. Bar, 10 μm.

**Expanded View Figure 7:**
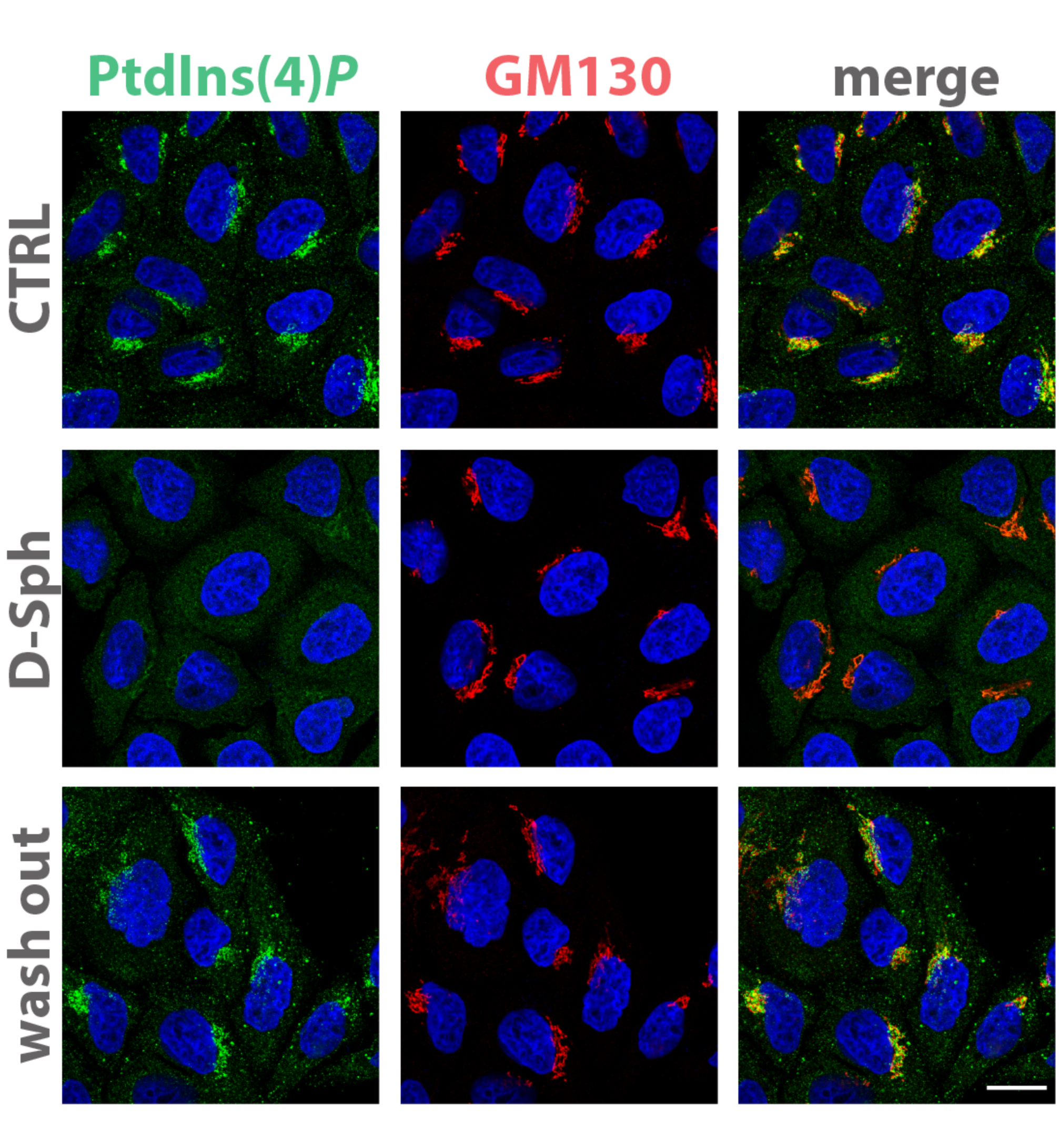
SL flow controls PtdIns(4)P levels at the Golgi. Cells treated either with vehicle (EtOH), D-Sph (30μM) for 30 min, or treated with D-Sph (30μM) for 30 min and washed out for 4 hours were fixed and permeabilized as in **(Figure 3A)** and stained with DAPI (blue), an anti-GM130 antibody (red) and anti PtdIns(4)*P-*antibody (green). Bar, 10 μm.

**Expanded View Figure 8:**
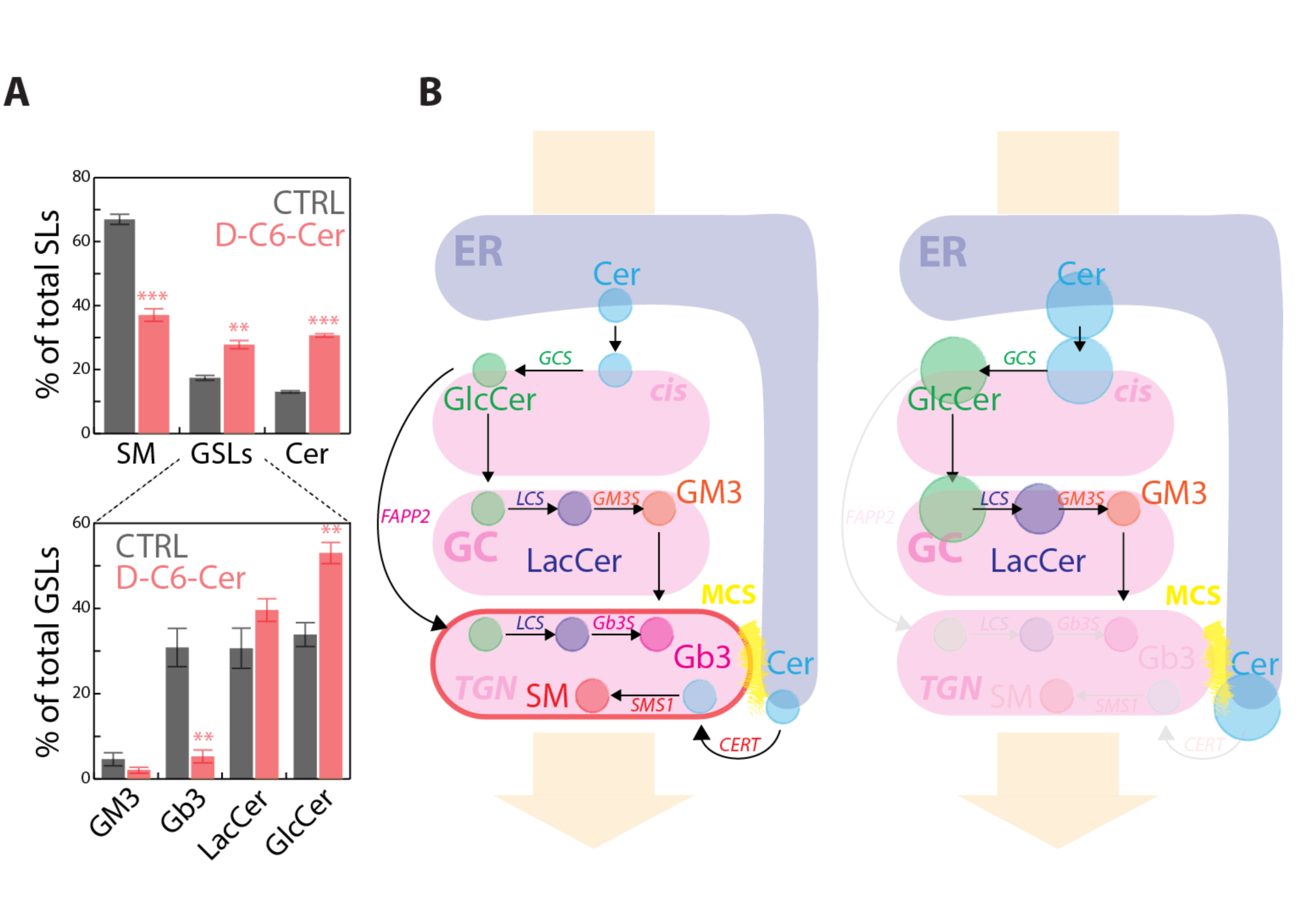
Sustained SL flow inhibits GSLs synthesis at the TGN. **A)** Cells treated either with vehicle (EtOH), D-C6-Cer (10μM, 120) were pulse labelled with ^3^H-D-Sph for SL and GSL synthesis assessment. The percentage of total radioactivity associated with SM, Cer and GSLs (i.e., GlcCer, LacCer, Gb3, and GM3) in the different conditions was quantitated after lipid extraction and HPTLC separation (upper panel). The percentage of GSLs radioactivity associated with GlcCer, LacCer, Gb3, and GM3 in the different conditions is also reported. Data are means ± S.E.M. of 4 independent experiments. **B)** Schematic representation of sustained SL flow on CERT and FAPP2 dependent SL synthetic pathways.

**Expanded View Figure 9:**
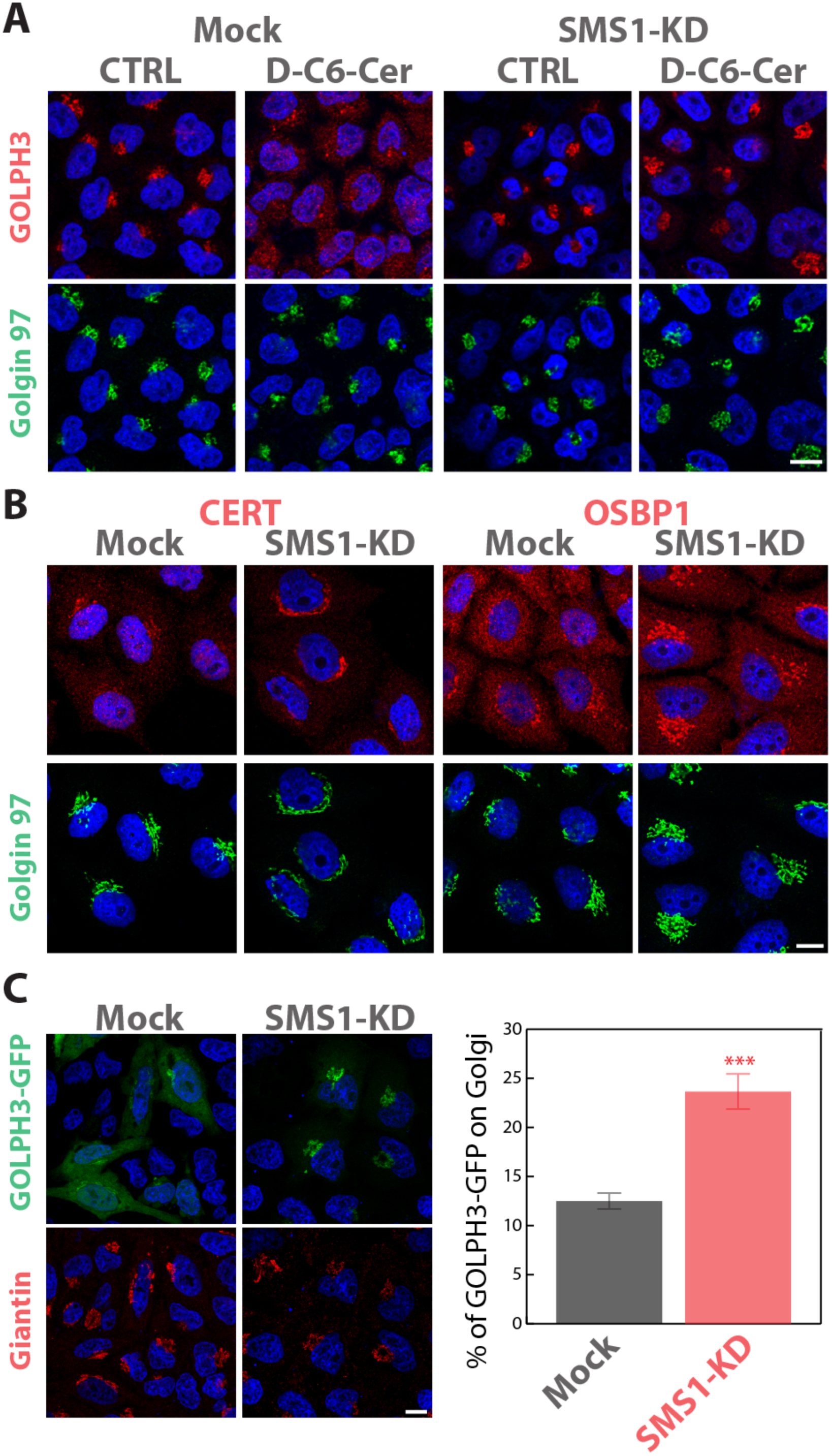
SM synthesis controls PtdIns(4)P binding proteins association to the TGN. **A)** Cells SMS1-KD were treated with either with EtOH (CTRL), or D-C6-Cer (10μM) for 2 hours, fixed, and stained with DAPI (blue), an anti-Golgin97 antibody (green) and anti-GOLPH3 antibody (red). **B)** Cells either mock treated or SMS1-KD were fixed, and stained with DAPI (blue), an anti-Golgin97 antibody (green) and either anti CERT or anti OSBP1 antibodies (red). **C)** Cells SMS1-KD were transfected with GFP- tagged GOLPH3 and fixed, and stained with DAPI (blue), and anti-Giantin antibody (red). The percentage of GOLPH3-GFP associated with the Golgi is reported in mock interfered (n=27) and SMS1-KD cells (n=27). Data are means ± S.E.M.

**Expanded View Figure 10:**
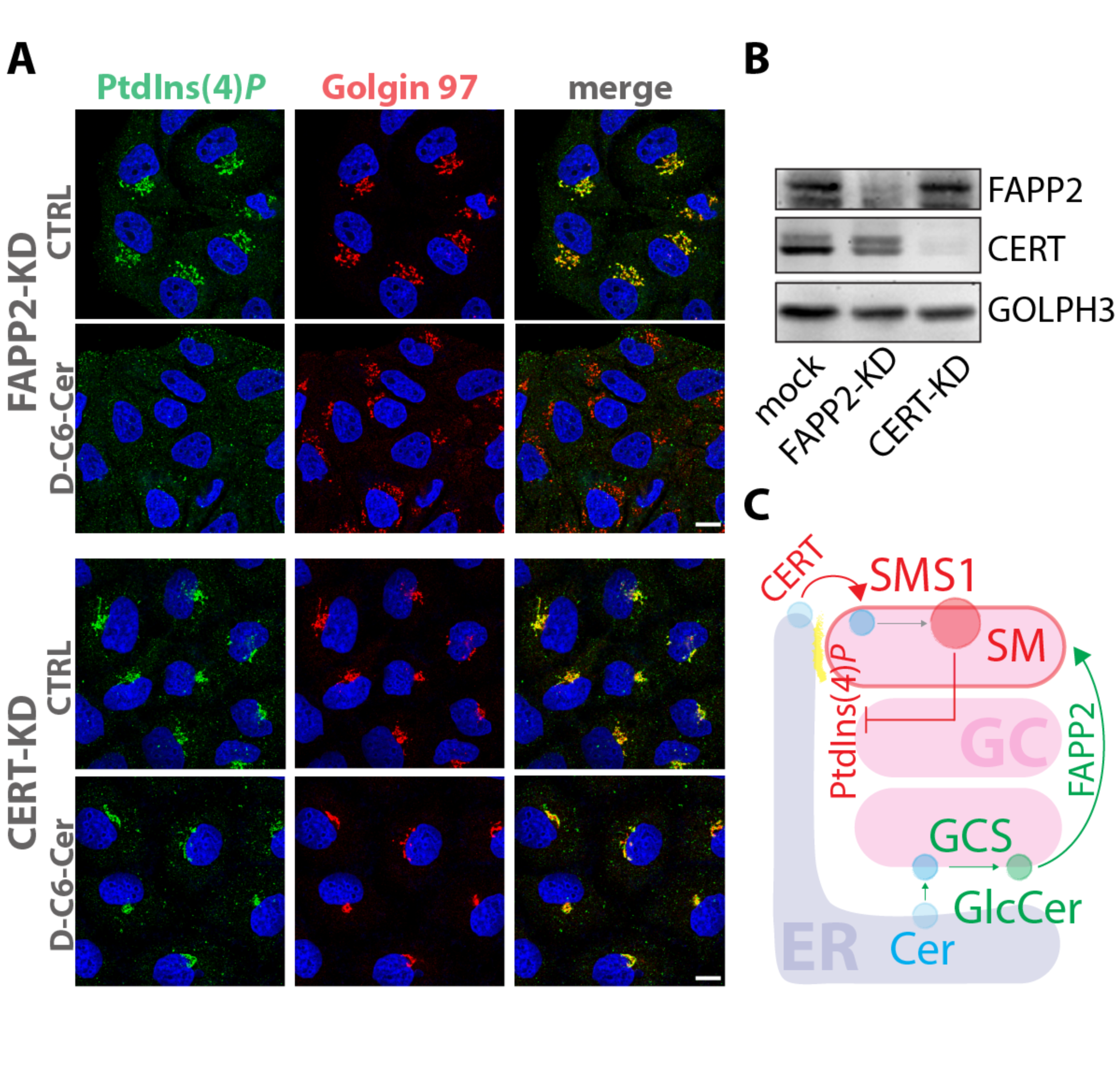
The CERT-SMS1 axis controls PtdIns(4)P levels at the TGN. **A)** Cells treated with FAPP2 or CERT directed siRNAs were treated with either with EtOH, or D-C6-Cer (10μM) for 30 min, fixed, permeabilized and stained with DAPI (blue), an anti-Golgin97 antibody (red) and a specific anti PtdIns(4)*P*-antibody (green; left panels). Bars, 10 μm. **B)** FAPP2 and CERT KD efficiencies were evaluated by Western Blotting. **C)** Schematic representation of CERT-SMS1 axis control on TGN PtdIns(4)*P*.

**Expanded View Figure 12:**
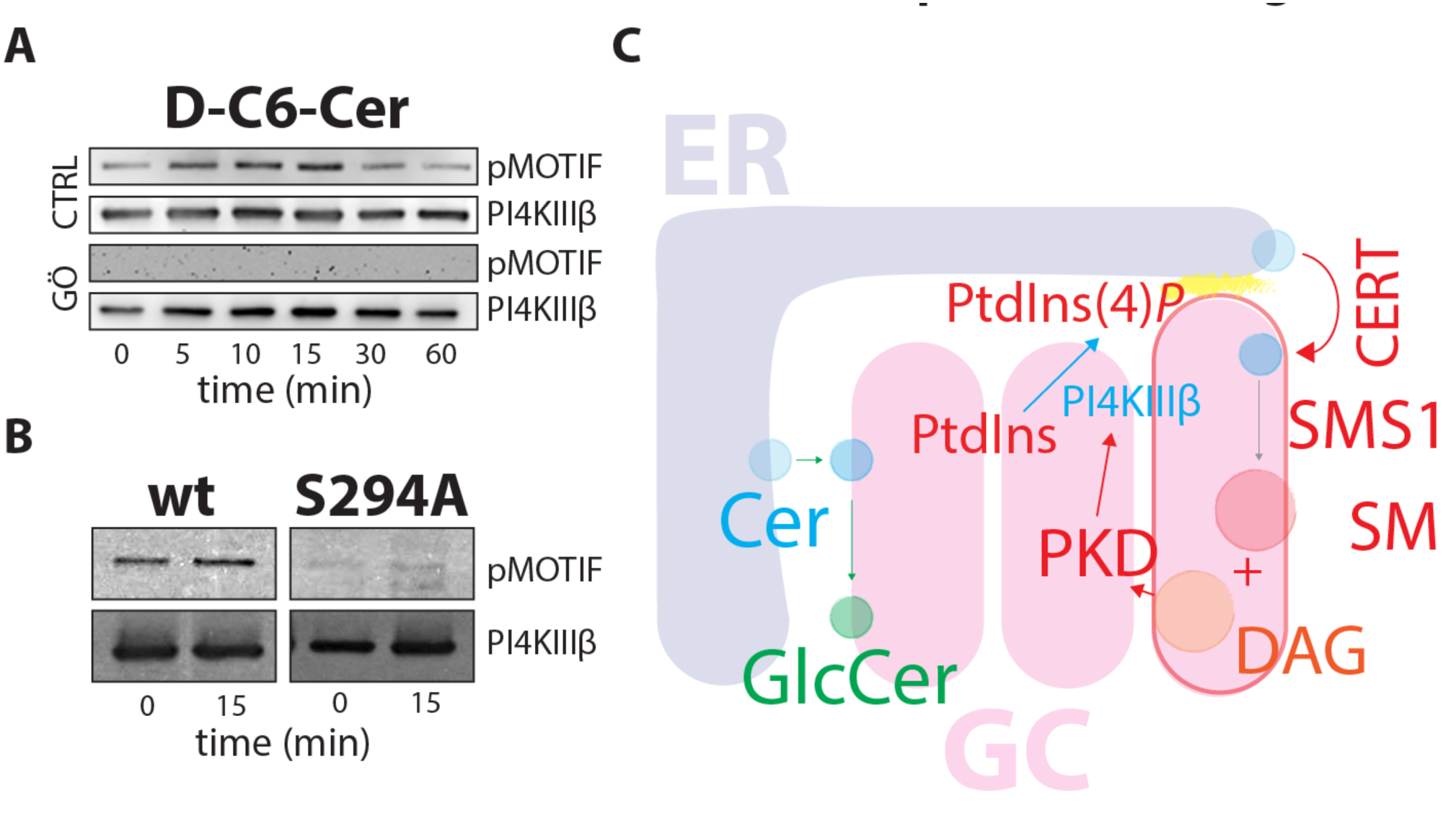
PI4KIIIβ response to SL flow. **A)** Cells expressing PI4KIIIβ-GFP were treated for the indicated times with D-C6-Cer (10μM) lysed, and protein lysates were subjected to immunoprecipitation by the use of anti-GFP antibody, SDS-PAGE and Western Blotting. PKD dependent phosphorylation of PI4KIIIβ was revealed by pMOTIF antibody and compared to the amount of PI4KIIIβ (detected by anti-PI4KIIIβ antibody) immunoprecipitated at each time point. The PKD inhibitor Gö 6976 (10 μM, pre treatment 1 hour) was used as a specificity control. **B)** Cells expressing PI4KIIIβ-GFP-wt (wt) or PI4KIIIβ-GFP-S294A mutant (S294A) were treated for the indicated times with D-C6-Cer (10μM) lysed, and protein lysates were subjected to immunoprecipitation by the use of anti-GFP antibody, SDS-PAGE and Western Blotting. PKD dependent phosphorylation of PI4KIIIβ was revealed by pMOTIF antibody and compared to the amount of PI4KIIIβ (detected by anti-PI4KIIIβ antibody) immunoprecipitated at each time point. **C)** Schematic representation of SL flow mediated phosphorylation of PI4KIIIβ.

**Expanded View Figure 13:**
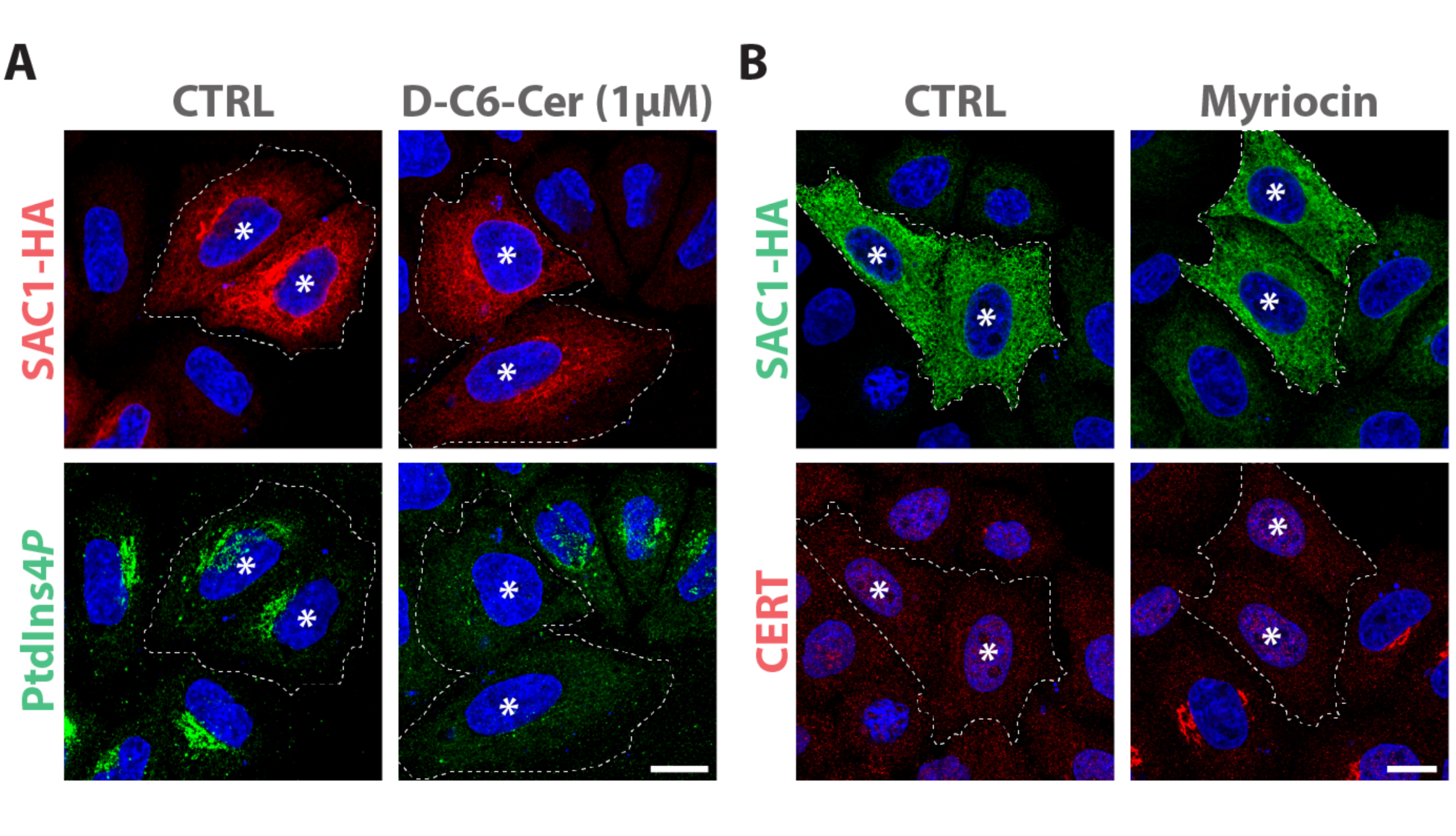
Sac1 consumes PtdIns(4)P in response to sustained SL flow. **A)** Cells expressing HA-SAC1 (red) were treated with D-C6-Cer (1μM, 1 hour) and the PtdIns(4)*P* (green) levels at the TGN were evaluated. **B)** Cells expressing HA-SAC1 (green) were treated with myriocin (2,5μM, 24 hours) and CERT (red) recruitment at the TGN was evaluated. Data are representative of at least three independent experiments. Bars, 10 μm.

**Expanded View Figure 14:**
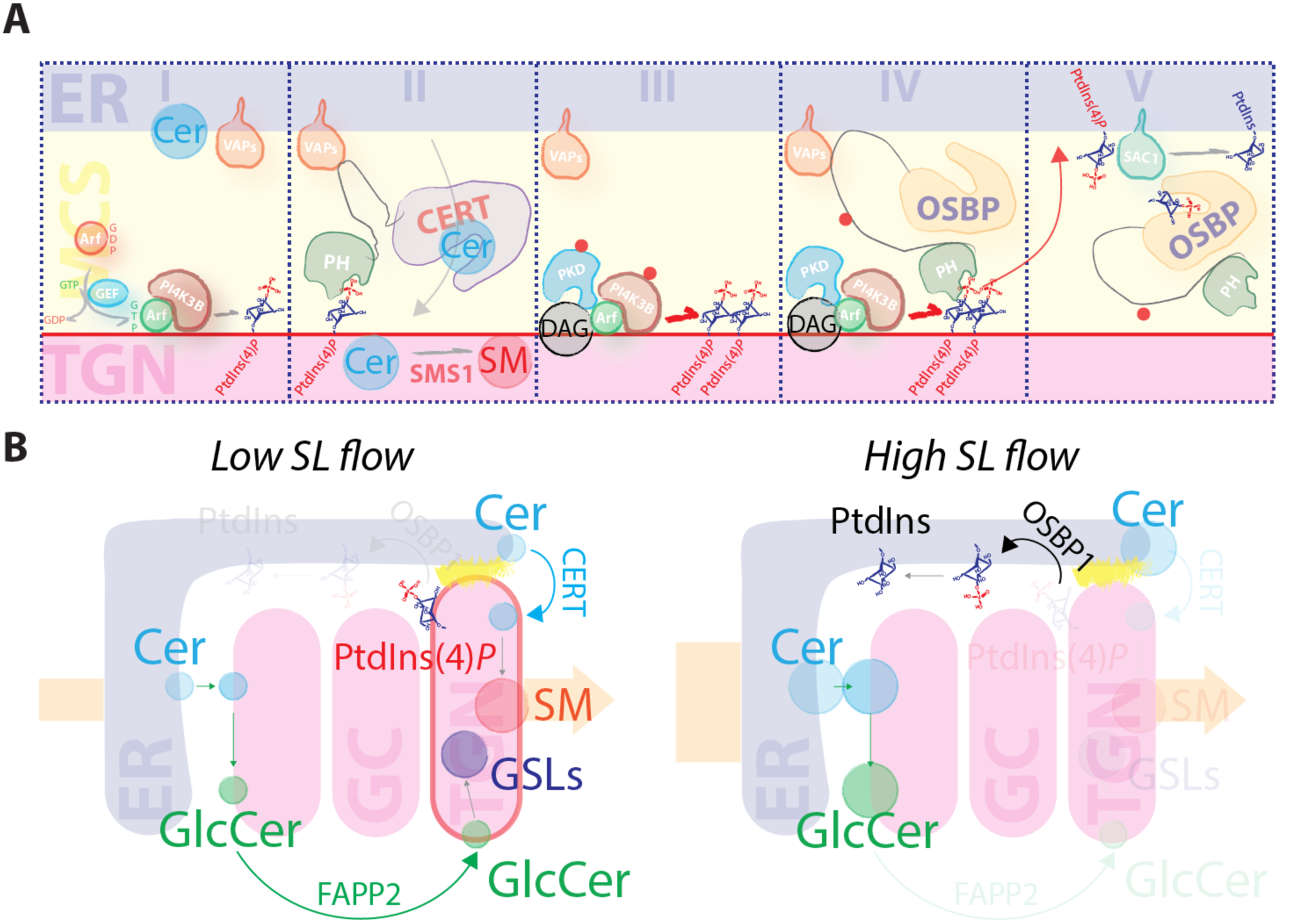
SL flow to the Golgi controls phosphoinositide turnover at the TGN. A) Schematic representation of SL controlled phosphoinositide turnover at the ER-TGN MCSs. **Phase I:** ARF1 recruits PI4KIIIβ to the TGN to produce PtdIns(4)*P*. **Phase II:** CERT transfers Cer from the ER to the TGN by contacting VAPs in the ER and PtdIns(4)*P* on the TGN. CERT transferred Cer is converted to SM by SMS1. **Phase III:** SMS1 produced DAG recruits (together with ARF) and locally activates PKD. PKD phosphorylates both PI4KIIIβ to increase PtdIns(4)*P* production. **Phase IV:** OSBP1 is recruited by PtdIns(4)*P.* PKD-phosphorylates OSBP1. **Phase IV:** OSBP1 transfers PtdIns(4)*P* to the ER for its dephosphorylation operated by SAC1. **B)** Schematic representation of TGN response to sustained SL flow. At low SL flow (left scheme) Cer and GlcCer are efficiently transported to the TGN by the PtdIns(4)*P* binding proteins CERT and FAPP2, respectively to yield SM and complex GSLs. At high SL flow (right scheme) PtdIns(4)*P* is transported to the ER by OSBP1 and dephosphorylated to PtdIns. As a consequence CERT and FAPP2 cannot deliver further SL precursors (Cer and GlcCer) to the TGN resulting in reduced production of SM and complex GSLs.

**Figure 4- Figure supplement 2:**
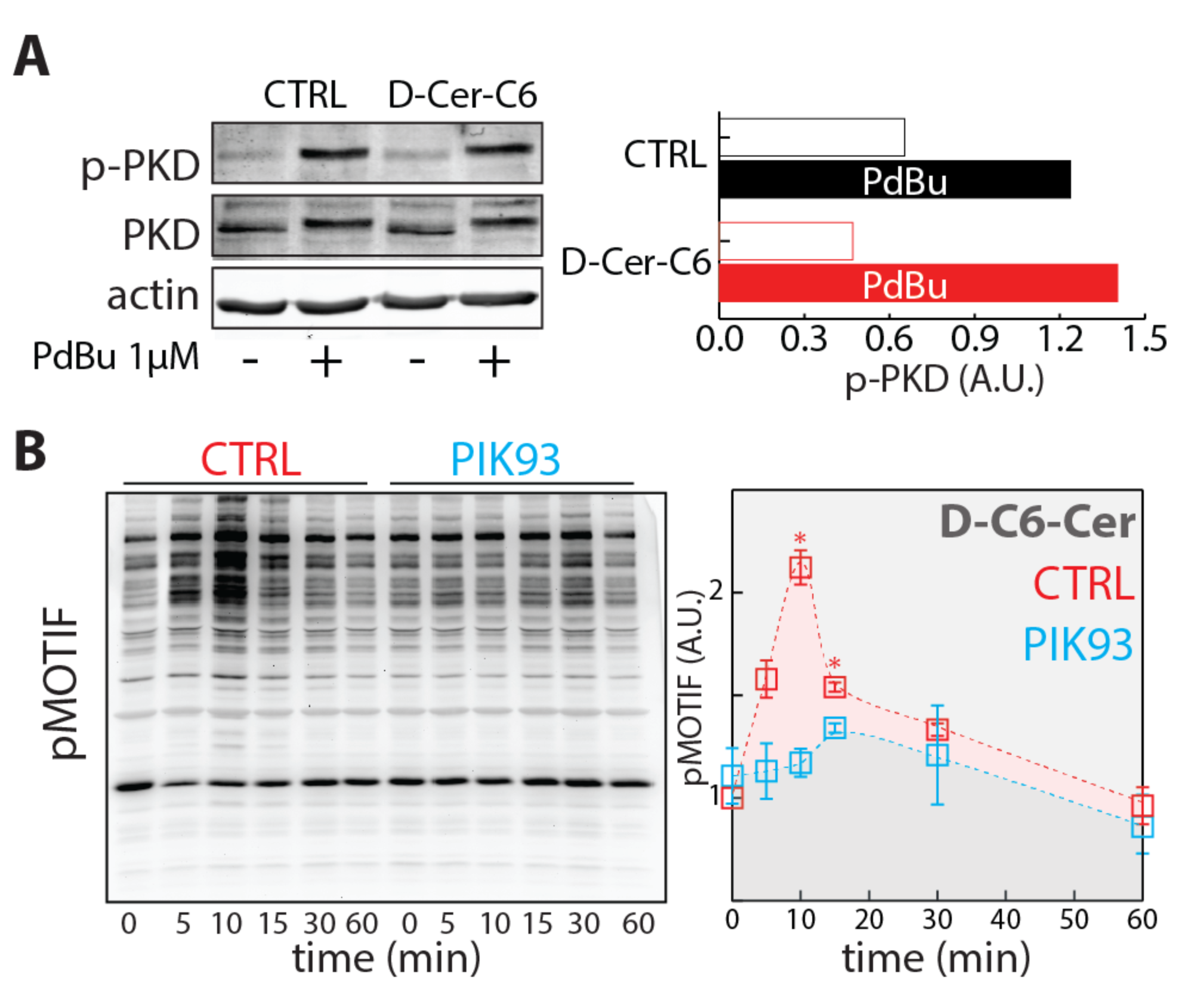
SL induced PKD activation requires PtdIns(4)P. **A)** Cells pre-treated with vehicle (EtOH) or 10μM D-C6-Cer (10 μM) for 30 min were treated for 15 min with PdBu (1μM) lysed and lysates were processed for SDS-PAGE and Western Blotting. PKD activation expressed as normalized p-PKD signal is reported. **B)** HeLa cells pre-treated with vehicle (EtOH) or PIK93 (250nM) for 15 min were subjected to D-C6-Cer (10 μM) treatment for the indicated times, lysed and lysates were processed for SDS-PAGE and Western Blotting. pMOTIF antibody was used to monitor PKD activation (left panel). PKD activation in both control (CTRL; red) and PIK93 (cyan) treated cells was evaluated by quantitating phosphorylation of PKD substrates during D-C6-Cer (10μM) administration. Data are means of at least 3 independent experiments ± standard deviation.

## Bibliography

Bajaj Pahuja K, Wang J, Blagoveshchenskaya A, Lim L, Madhusudhan MS, Mayinger P, Schekman R (2015) Phosphoregulatory protein 14-3-3 facilitates SAC1 transport from the endoplasmic reticulum. Proceedings of the National Academy of Sciences of the United States of America 112: E3199–206

Blagoveshchenskaya A, Cheong FY, Rohde HM, Glover G, Knodler A, Nicolson T, Boehmelt G, Mayinger P (2008) Integration of Golgi trafficking and growth factor signaling by the lipid phosphatase SAC1. The Journal of cell biology 180: 803–12

Breslow DK, Collins SR, Bodenmiller B, Aebersold R, Simons K, Shevchenko A, Ejsing CS, Weissman JS (2010) Orm family proteins mediate sphingolipid homeostasis. Nature 463: 1048–53

Breslow DK, Weissman JS (2010) Membranes in balance: mechanisms of sphingolipid homeostasis. Molecular cell 40: 267–79

Burd CG, Emr SD (1998) Phosphatidylinositol(3)-phosphate signaling mediated by specific binding to RING FYVE domains. Molecular cell 2: 157–62

Cantalupo A, Zhang Y, Kothiya M, Galvani S, Obinata H, Bucci M, Giordano FJ, Jiang XC, Hla T, Di Lorenzo A (2015) Nogo-B regulates endothelial sphingolipid homeostasis to control vascular function and blood pressure. Nature medicine 21: 1028–37

Carpenter AE, Jones TR, Lamprecht MR, Clarke C, Kang IH, Friman O, Guertin DA, Chang JH, Lindquist RA, Moffat J, Golland P, Sabatini DM (2006) CellProfiler: image analysis software for identifying and quantifying cell phenotypes. Genome biology 7: R100

Chen J, Deng F, Li J, Wang QJ (2008) Selective binding of phorbol esters and diacylglycerol by individual C1 domains of the PKD family. The Biochemical journal 411: 333–42

Chung J, Torta F, Masai K, Lucast L, Czapla H, Tanner LB, Narayanaswamy P, Wenk MR, Nakatsu F, De Camilli P (2015) INTRACELLULAR TRANSPORT. PI4P/phosphatidylserine countertransport at ORP5- and ORP8-mediated ER-plasma membrane contacts. Science 349: 428–32

Cruz-Garcia D, Ortega-Bellido M, Scarpa M, Villeneuve J, Jovic M, Porzner M, Balla T, Seufferlein T, Malhotra V (2013) Recruitment of arfaptins to the trans-Golgi network by PI(4)P and their involvement in cargo export. The EMBO journal 32: 1717–29

D‘Angelo G, Polishchuk E, Di Tullio G, Santoro M, Di Campli A, Godi A, West G, Bielawski J, Chuang CC, van der Spoel AC, Platt FM, Hannun YA, Polishchuk R, Mattjus P, De Matteis MA (2007) Glycosphingolipid synthesis requires FAPP2 transfer of glucosylceramide. Nature 449: 62–7

D‘Angelo G, Uemura T, Chuang CC, Polishchuk E, Santoro M, Ohvo-Rekila H, Sato T, Di Tullio G, Varriale A, D‘Auria S, Daniele T, Capuani F, Johannes L, Mattjus P, Monti M, Pucci P, Williams RL, Burke JE, Platt FM, Harada A et al. (2013) Vesicular and non-vesicular transport feed distinct glycosylation pathways in the Golgi. Nature 501: 116–20

D‘Angelo G, Vicinanza M, De Matteis MA (2008) Lipid-transfer proteins in biosynthetic pathways. Current opinion in cell biology 20: 360–70

De Matteis MA, Luini A (2008) Exiting the Golgi complex. Nature reviews Molecular cell biology 9: 273–84

De Matteis MA, Rega LR (2015) Endoplasmic reticulum-Golgi complex membrane contact sites. Current opinion in cell biology 35: 43–50

De Matteis MA, Wilson C, D‘Angelo G (2013) Phosphatidylinositol-4-phosphate: the Golgi and beyond. BioEssays: news and reviews in molecular, cellular and developmental biology 35: 612–22

Deng Y, Rivera-Molina FE, Toomre DK, Burd CG (2016) Sphingomyelin is sorted at the trans Golgi network into a distinct class of secretory vesicle. Proceedings of the National Academy of Sciences of the United States of America 113: 6677–82

Dippold HC, Ng MM, Farber-Katz SE, Lee SK, Kerr ML, Peterman MC, Sim R, Wiharto PA, Galbraith KA, Madhavarapu S, Fuchs GJ, Meerloo T, Farquhar MG, Zhou H, Field SJ (2009) GOLPH3 bridges phosphatidylinositol-4- phosphate and actomyosin to stretch and shape the Golgi to promote budding. Cell 139: 337–51

Doppler H, Storz P, Li J, Comb MJ, Toker A (2005) A phosphorylation state-specific antibody recognizes Hsp27, a novel substrate of protein kinase D. The Journal of biological chemistry 280: 15013–9

Dowler S, Currie RA, Campbell DG, Deak M, Kular G, Downes CP, Alessi DR (2000) Identification of pleckstrin-homology-domain-containing proteins with novel phosphoinositide-binding specificities. The Biochemical journal 351: 19–31

Duran JM, Campelo F, van Galen J, Sachsenheimer T, Sot J, Egorov MV, Rentero C, Enrich C, Polishchuk RS, Goni FM, Brugger B, Wieland F, Malhotra V (2012) Sphingomyelin organization is required for vesicle biogenesis at the Golgi complex. The EMBO journal 31: 4535–46

Fugmann T, Hausser A, Schoffler P, Schmid S, Pfizenmaier K, Olayioye MA (2007) Regulation of secretory transport by protein kinase D-mediated phosphorylation of the ceramide transfer protein. The Journal of cell biology 178: 15–22

Godi A, Di Campli A, Konstantakopoulos A, Di Tullio G, Alessi DR, Kular GS, Daniele T, Marra P, Lucocq JM, De Matteis MA (2004) FAPPs control Golgi-to-cell-surface membrane traffic by binding to ARF and PtdIns(4)P. Nature cell biology 6: 393–404

Hammond GR, Schiavo G, Irvine RF (2009) Immunocytochemical techniques reveal multiple, distinct cellular pools of PtdIns4P and PtdIns(4,5)P(2). The Biochemical journal 422: 23–35

Hanada K, Kumagai K, Yasuda S, Miura Y, Kawano M, Fukasawa M, Nishijima M (2003) Molecular machinery for non-vesicular trafficking of ceramide. Nature 426: 803–9

Hannun YA, Obeid LM (2008) Principles of bioactive lipid signalling: lessons from sphingolipids. Nature reviews Molecular cell biology 9: 139–50

Hausser A, Storz P, Martens S, Link G, Toker A, Pfizenmaier K (2005) Protein kinase D regulates vesicular transport by phosphorylating and activating phosphatidylinositol-4 kinase IIIbeta at the Golgi complex. Nature cell biology 7: 880–6

He J, Scott JL, Heroux A, Roy S, Lenoir M, Overduin M, Stahelin RV, Kutateladze TG (2011) Molecular basis of phosphatidylinositol 4-phosphate and ARF1 GTPase recognition by the FAPP1 pleckstrin homology (PH) domain. The Journal of biological chemistry 286: 18650–7

Holthuis JC, Menon AK (2014) Lipid landscapes and pipelines in membrane homeostasis. Nature 510: 48–57

Horvath A, Sutterlin C, Manning-Krieg U, Movva NR, Riezman H (1994) Ceramide synthesis enhances transport of GPI-anchored proteins to the Golgi apparatus in yeast. The EMBO journal 13: 3687–95

Hou Q, Jin J, Zhou H, Novgorodov SA, Bielawska A, Szulc ZM, Hannun YA, Obeid LM, Hsu YT (2011) Mitochondrially targeted ceramides preferentially promote autophagy, retard cell growth, and induce apoptosis. Journal of lipid research 52: 278–88

Huitema K, van den Dikkenberg J, Brouwers JF, Holthuis JC (2004) Identification of a family of animal sphingomyelin synthases. The EMBO journal 23: 33–44

Ichikawa S, Ozawa K, Hirabayashi Y (1998) Molecular cloning and characterization of the mouse ceramide glucosyltransferase gene. Biochemical and biophysical research communications 253: 707–11

Klemm RW, Ejsing CS, Surma MA, Kaiser HJ, Gerl MJ, Sampaio JL, de Robillard Q, Ferguson C, Proszynski TJ, Shevchenko A, Simons K (2009) Segregation of sphingolipids and sterols during formation of secretory vesicles at the trans-Golgi network. The Journal of cell biology 185: 601–12

Kumagai K, Kawano M, Shinkai-Ouchi F, Nishijima M, Hanada K (2007) Interorganelle trafficking of ceramide is regulated by phosphorylation-dependent cooperativity between the PH and START domains of CERT. The Journal of biological chemistry 282: 17758–66

Lemmon MA, Ferguson KM, O‘Brien R, Sigler PB, Schlessinger J (1995) Specific and high-affinity binding of inositol phosphates to an isolated pleckstrin homology domain. Proceedings of the National Academy of Sciences of the United States of America 92: 10472–6

Lenoir M, Coskun U, Grzybek M, Cao X, Buschhorn SB, James J, Simons K, Overduin M (2010) Structural basis of wedging the Golgi membrane by FAPP pleckstrin homology domains. EMBO reports 11: 279–84

Liljedahl M, Maeda Y, Colanzi A, Ayala I, Van Lint J, Malhotra V (2001) Protein kinase D regulates the fission of cell surface destined transport carriers from the trans-Golgi network. Cell 104: 409–20

Lingwood D, Simons K (2010) Lipid rafts as a membrane-organizing principle. Science 327: 46–50

Lippincott-Schwartz J, Yuan LC, Bonifacino JS, Klausner RD (1989) Rapid redistribution of Golgi proteins into the ER in cells treated with brefeldin A: evidence for membrane cycling from Golgi to ER. Cell 56: 801–13

Malhotra V, Campelo F (2011) PKD regulates membrane fission to generate TGN to cell surface transport carriers. Cold Spring Harbor perspectives in biology 3

Mesmin B, Bigay J, Moser von Filseck J, Lacas-Gervais S, Drin G, Antonny B (2013) A four-step cycle driven by PI(4)P hydrolysis directs sterol/PI(4)P exchange by the ER-Golgi tether OSBP. Cell 155: 830–43

Morad SA, Cabot MC (2013) Ceramide-orchestrated signalling in cancer cells. Nat Rev Cancer 13: 51–65

Nakano A, Luini A (2010) Passage through the Golgi. Current opinion in cell biology 22: 471–8

Nhek S, Ngo M, Yang X, Ng MM, Field SJ, Asara JM, Ridgway ND, Toker A (2010) Regulation of oxysterol-binding protein Golgi localization through protein kinase D-mediated phosphorylation. Molecular biology of the cell 21: 2327–37

Ogretmen B, Pettus BJ, Rossi MJ, Wood R, Usta J, Szulc Z, Bielawska A, Obeid LM, Hannun YA (2002) Biochemical mechanisms of the generation of endogenous long chain ceramide in response to exogenous short chain ceramide in the A549 human lung adenocarcinoma cell line. Role for endogenous ceramide in mediating the action of exogenous ceramide. The Journal of biological chemistry 277: 12960–9

Pagliuso A, Valente C, Giordano LL, Filograna A, Li G, Circolo D, Turacchio G, Marzullo VM, Mandrich L, Zhukovsky MA, Formiggini F, Polishchuk RS, Corda D, Luini A (2016) Golgi membrane fission requires the CtBP1-S/BARS-induced activation of lysophosphatidic acid acyltransferase delta. Nature communications 7: 12148

Patterson GH, Hirschberg K, Polishchuk RS, Gerlich D, Phair RD, Lippincott-Schwartz J (2008) Transport through the Golgi apparatus by rapid partitioning within a two-phase membrane system. Cell 133: 1055–67

Rykx A, De Kimpe L, Mikhalap S, Vantus T, Seufferlein T, Vandenheede JR, Van Lint J (2003) Protein kinase D: a family affair. FEBS letters 546: 81–6

Schneider CA, Rasband WS, Eliceiri KW (2012) NIH Image to ImageJ: 25 years of image analysis. Nature methods 9: 671–5

Sharpe HJ, Stevens TJ, Munro S (2010) A comprehensive comparison of transmembrane domains reveals organelle-specific properties. Cell 142: 158–69

Siow DL, Wattenberg BW (2012) Mammalian ORMDL proteins mediate the feedback response in ceramide biosynthesis. The Journal of biological chemistry 287: 40198–204

Stauffer TP, Ahn S, Meyer T (1998) Receptor-induced transient reduction in plasma membrane PtdIns(4,5)P2 concentration monitored in living cells. Curr Biol 8: 343–6

Subathra M, Qureshi A, Luberto C (2011) Sphingomyelin synthases regulate protein trafficking and secretion. PloS one 6: e23644

Thomaseth C, Weber P, Hamm T, Kashima K, Radde N (2013) Modeling sphingomyelin synthase 1 driven reaction at the Golgi apparatus can explain data by inclusion of a positive feedback mechanism. Journal of theoretical biology 337: 174–80

Toth B, Balla A, Ma H, Knight ZA, Shokat KM, Balla T (2006) Phosphatidylinositol 4-kinase IIIbeta regulates the transport of ceramide between the endoplasmic reticulum and Golgi. The Journal of biological chemistry 281: 36369–77

Vacaru AM, Tafesse FG, Ternes P, Kondylis V, Hermansson M, Brouwers JF, Somerharju P, Rabouille C, Holthuis JC (2009) Sphingomyelin synthase-related protein SMSr controls ceramide homeostasis in the ER. The Journal of cell biology 185: 1013–27

Valente C, Turacchio G, Mariggio S, Pagliuso A, Gaibisso R, Di Tullio G, Santoro M, Formiggini F, Spano S, Piccini D, Polishchuk RS, Colanzi A, Luini A, Corda D (2012) A 14-3-3gamma dimer-based scaffold bridges CtBP1-S/BARS to PI(4)KIIIbeta to regulate post-Golgi carrier formation. Nature cell biology 14: 343–54

van Galen J, Campelo F, Martinez-Alonso E, Scarpa M, Martinez-Menarguez JA, Malhotra V (2014) Sphingomyelin homeostasis is required to form functional enzymatic domains at the trans-Golgi network. The Journal of cell biology 206: 609–18

van Meer G, Voelker DR, Feigenson GW (2008) Membrane lipids: where they are and how they behave. Nature reviews Molecular cell biology 9: 112–24

Varnai P, Balla T (1998) Visualization of phosphoinositides that bind pleckstrin homology domains: calcium- and agonist-induced dynamic changes and relationship to myo-[3H]inositol-labeled phosphoinositide pools. The Journal of cell biology 143: 501–10

Villani M, Subathra M, Im YB, Choi Y, Signorelli P, Del Poeta M, Luberto C (2008) Sphingomyelin synthases regulate production of diacylglycerol at the Golgi. The Biochemical journal 414: 31–41

Wakana Y, Kotake R, Oyama N, Murate M, Kobayashi T, Arasaki K, Inoue H, Tagaya M (2015) CARTS biogenesis requires VAP-lipid transfer protein complexes functioning at the endoplasmic reticulum-Golgi interface. Molecular biology of the cell 26: 4686–99

Wang E, Norred WP, Bacon CW, Riley RT, Merrill AH, Jr. (1991) Inhibition of sphingolipid biosynthesis by fumonisins. Implications for diseases associated with Fusarium moniliforme. The Journal of biological chemistry 266: 14486–90

Wang YJ, Wang J, Sun HQ, Martinez M, Sun YX, Macia E, Kirchhausen T, Albanesi JP, Roth MG, Yin HL (2003) Phosphatidylinositol 4 phosphate regulates targeting of clathrin adaptor AP-1 complexes to the Golgi. Cell 114: 299–310

Weber P, Hornjik M, Olayioye MA, Hausser A, Radde NE (2015) A computational model of PKD and CERT interactions at the trans-Golgi network of mammalian cells. BMC systems biology 9: 9

Yamaji T, Hanada K (2015) Sphingolipid metabolism and interorganellar transport: localization of sphingolipid enzymes and lipid transfer proteins. Traffic 16: 101–22

